# Synchronization of the segmentation clock using synthetic cell-cell signaling

**DOI:** 10.1101/2024.11.04.617523

**Authors:** Akihiro Isomura, Daisuke Asanuma, Ryoichiro Kageyama

## Abstract

Tight coordination of cell-cell signaling in space and time is vital for self-organization in tissue patterning. During vertebrate development, the segmentation clock drives oscillatory gene expression in the presomitic mesoderm (PSM), leading to the periodic formation of somites. Oscillatory gene expression is synchronized at the cell population level; inhibition of Delta-Notch signaling results in the loss of synchrony and the fusion of somites. However, it remains unclear how cell-cell signaling couples oscillatory gene expression and controls synchronization. Here, we report the reconstitution of synchronized oscillation in PSM organoids by synthetic cell-cell signaling with designed ligand-receptor pairs. Optogenetic assays uncovered that the intracellular domains of synthetic ligands play key roles in dynamic cell-cell communication. Oscillatory coupling using synthetic cell-cell signaling recovered the synchronized oscillation in PSM cells deficient for Delta-Notch signaling; non-oscillatory coupling did not induce recovery. This study reveals the mechanism by which ligand-receptor molecules coordinate the synchronization of the segmentation clock, and provides direct evidence of oscillatory cell-cell communication in the segmentation clock.

## Introduction

Cell-cell signaling provides the means to send and receive information that is essential for embryonic development and tissue homeostasis. In many cases, cells relay the information carried by signaling molecules, such as morphogens; however, stochastic biochemical reactions and molecular diffusion can reduce the amount of information and thereby prevent reliable communication between distant cells at the tissue level^1,2^. This raises the fundamental questions of whether and how cells communicate precise information to each other in space and time.

One solution to this communication problem is synchronization of pulsatile or oscillatory cell-cell signaling, in which cell-cell communication coordinates simultaneous signaling activation at the cell population level and mitigates fluctuations between individual cells. Biofilms of *Bacillus subtilis* cells display collective oscillations of growth, metabolic activity and gene expression, which support cell survival^3,4^. During the aggregation stage of the social amoeba *Dictyostelium*, cells produce a chemoattractant cAMP signal in a pulsatile manner, which is relayed and propagated as a traveling wave that induces collective cell movement^5^. *Drosophila* embryos utilize mitotic oscillatory waves for the proper scheduling of the body plan^6,7^. In vertebrates, a traveling wave of pulsatile Erk activity is involved in epidermal wound healing and osteoblast regeneration^8,9^. These examples highlight the relay of pulsatile signaling activities over sub-millimeter scales in diverse species; however, it is unclear how signaling molecules encode and convey oscillatory information in cell-cell communication.

The segmentation clock has a vital role in the control of morphogenesis via synchronized oscillatory gene expression^10,11^. During development, presomitic mesoderm (PSM) tissue of vertebrate embryos is periodically segmented into metameric structures, somites, which give rise to bones and muscles. In a PSM cell population, the timing of somitogenesis is controlled by the segmentation clock through synchronization of gene expression. A transcriptional repressor, Hes7 in mice and Her1/7 in zebrafish, is an effector gene of Delta-Notch signaling, and forms a negative feedback loop by repressing its own transcription, thereby driving oscillatory gene expression in the segmentation clock^10,11^. Deficiency of Notch signaling causes *Hes7* oscillation to diminish at the tissue level and results in severe defects in somite structures^12–14^. Isolated single PSM cells can maintain *Hes7* oscillation in the absence of cell-cell contact, but the phases and periods are variable. By contrast, re-aggregation of dissociated PSM cells reconstitutes synchronized oscillation and produces uniform phases and periods^15–17^. Inhibition of Notch signaling blocks the recovery of synchronization in re-aggregated PSM cells^15,16^, indicating that Notch signaling is a cell-cell coupling mechanism for the synchronized oscillation of the segmentation clock. In zebrafish, *Her1*/*7* directly controls promoter activity of the Notch ligand *DeltaC*, while DeltaC ligands transactivate *Her1*/*7* expression in neighboring cells^11,18–20^. A minimal model based on these observations suggested that the core oscillator gene *Hes7* or *Her1*/*7* drives oscillatory Delta expression, which then transmits core oscillator information to neighboring cells for synchronization^11,18^. Nevertheless, evidence that oscillatory cell-cell signaling couples PSM cells and further initiates self-organization of synchronization is still absent.

Direct testing of the minimal cell-cell coupling model for synchronization has been a challenge due to the complexity of Notch signaling. The functional roles of the extracellular domains of Delta-Notch molecules in cell-cell signaling has been investigated^21,22^; however, their regulatory roles in the segmentation clock remain obscure. In mouse PSM, the *lunatic fringe* (*Lfng*) gene modulates Notch activity by glycosylation of Delta and Notch extracellular domains^23,24^ and is additionally required for synchronization of Hes7 oscillations^17,25–27;^ the details of these actions are still unresolved. Zebrafish embryos with mutation of the Notch ligand *DeltaD* show loss of synchrony in *Her1*/*7* oscillations; as the *DeltaD* expression pattern *per se* is not oscillatory in wild-type embryos, the functional role of *DeltaD* in synchronization is unclear^28,29^. These observations are not consistent with the minimal model in which Hes7 or Her1/7 oscillators are coupled to each other by simple ligand-receptor interactions of Delta-Notch signaling.

One possible means to further explore the minimal model may be to replace the extracellular domains of Delta-Notch molecules with synthetic alternatives; this would reduce the complexity of the segmentation clock and thereby permit elucidation of the minimal design for cell-cell coupling and synchronization. Synthetic biology technologies offer an opportunity to test the functional capability of a hypothetical minimal design with customized genetic modules that could replace their natural counterparts^30,31^. Here, we investigate the importance of cell-cell coupling on the synchronization of the segmentation clock using a synthetic biology approach. We develop synthetic cell-cell signaling pathways that can transmit oscillatory information, and demonstrate that reconstitution of the synchronization of a segmentation clock can be achieved in organoids derived from mouse embryonic stem cells (mESCs).

### 1 Somitogenesis in organoids as a model system to test synthetic cell-cell signaling

We sought to establish a reliable system that could be used to test a range of minimal synthetic circuits with regard to synchronized oscillation of the segmentation clock. We previously developed an *in vitro* differentiation system that induced formation of PSM-like cells from the embryoid bodies of mESCs^32^. Other groups reported a protocol for PSM-like cell induction in gastruloid or trunk-like structures derived from mESCs; this protocol enables recapitulation of the periodic formation of somites with multiple tissue polarities^33,34^. By combining and modifying these protocols, we generated a new method that we refer to as mass-induced PSM (miPSM), which enables mass production of induced PSM-like cells and somite-like structures with a reduced number of handling steps and a smaller volume of culture medium (Extended Data Fig. 1A, supplementary text). We also constructed an mESC reporter line carrying an *Hes7*-promoter (pHes7) to monitor the segmentation clock in miPSM cells by fluorescence live-imaging, utilizing the fast maturing yellow fluorescent protein Achilles^17^ (Extended Data Fig. 1B). Time-lapse microscopy of organoid-cultures of miPSM derived from the pHes7 reporter mESC line displayed kinematic waves of pHes7 activity with about 2 h period that resembles somitogenesis in mouse embryos (Fig. 1, A to C). Analysis of the expression pattern of *Uncx4.1*, a marker for the caudal halves of somites, using *in situ* hybridization chain reaction (HCR), showed the presence of stripy patterns (Fig. 1D).

**Fig. 1.**
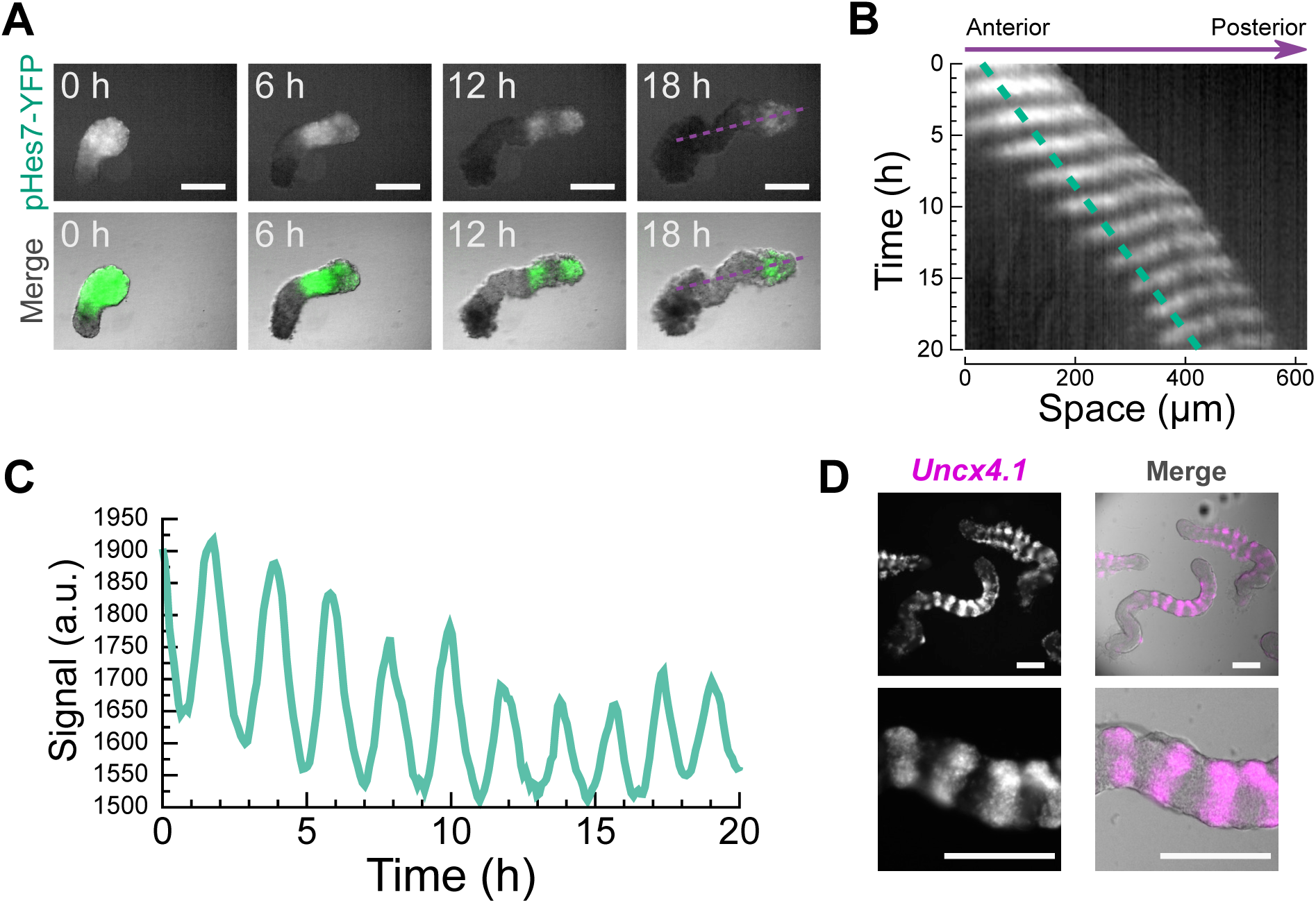
Organoids derived from miPSM cells as a model system to test synthetic cell-cell signaling. (A) Snapshots of yellow fluorescent protein (YFP; upper panels) and merged images of YFP and bright field channels (lower panels). Dashed lines (purple) indicate regions of interest (ROIs), in which signals were measured for a kymograph in (B). (B) A kymograph of organoids in an miPSM culture shows axis elongation and traveling waves of Hes7 signals. The dashed line (green) indicates the ROI analyzed in (C). (C) Oscillatory patterns of Hes7 reporter expression. (D) Hybridization chain reaction (HCR) v3.0 staining of *Uncx4.1* (somite marker gene). Scale bars, 200 *µ*m.

Mouse embryos that lack the *Delta-like1* (*Dll1*) gene show severely impaired somite structures^12,13^, indicating that Delta-Notch signaling plays an essential role in somitogenesis. We constructed *Dll1*-knockout (KO) mESC lines derived from pHes7 reporter mESCs and produced miPSM (Extended Data Fig. 2, A and B). We found that pHes7 reporter activity in *Dll1*-KO miPSM was weaker than in the wild-type (Extended Data Fig. 2C), as previously demonstrated in mouse *Dll1*-KO embryos^35^. The pHes7 reporter signals reduced after a few cycles of oscillation and continued to diminish thereafter, suggesting a loss of synchrony in the segmentation clock in *Dll1*-KO miPSM. Although axis elongation was apparent, in situ HCR analysis showed reduced *Uncx4.1* expression in *Dll1*-KO miPSM (Extended Data Fig. 2D). These data indicate that knockout of *Dll1* caused loss of synchronization in the segmentation clock and impairment of the somitogenesis program in miPSM, reminiscent of the phenotypes of *Dll1*-KO mouse embryos. Overall, these results demonstrate that the miPSM system resembles *in vivo* somitogenesis with its requirement for cell-cell communication via Delta-Notch signaling.

### 2 Optogenetic assays of the dynamic features of synthetic cell-cell signaling

Having established and validated the miPSM system, we next investigated potential candidates of synthetic signaling that could replace the native Delta-Notch signaling required for synchronization of the segmentation clock. Delta-Notch signaling has been suggested to be a dynamic communication channel: oscillatory activities of Notch signaling are transmitted from cells to cells during somitogenesis^10,11^. We made use of an optogenetic sender-receiver assay to identify and characterize synthetic cell-cell signaling pathways that are capable of transmitting oscillatory activity between cells^36^. Two types of cells were generated: photo-sensitive sender cells that triggered transcription of a ligand gene upon blue light illumination by the LightOn system^37^; and photo-insensitive receiver cells that carried an expression cassette of synthetic receptors. The expression cassette in the latter included a Tet activator and a reporter cassette of destabilized luciferase gene (*dLuc*) driven by a Tet-responsive element (TRE) promoter (Extended Data Fig. 3A). First, we determined whether the TRE promoter-driven luciferase reporter could detect dynamic cell-cell communication between sender and receiver cells. We generated photo-insensitive receiver cells carrying a Notch reporter system, in which the intracellular domain of mouse Notch1 was replaced with a Tet activator, and photo-sensitive sender cells producing natural ligand Delta-like1 (Dll1) upon blue light illumination (Extended Data Fig. 3, A and B). We applied cyclic blue light-stimulation with a 4-h period to a co-culture of sender and receiver cells. Time-lapse recording of bio-luminescence signals from the receiver cells showed periodic patterns with a 4-h period (Extended Data Fig. 3C), suggesting the successful transfer of oscillatory information from sender to receiver cells.

Next, we investigated synthetic cell-cell signaling patterns using a synNotch system^38^. In this system, the synNotch receptor (synNR) consists of a signal peptide derived from human CD8*α* for translocation to the cellular membrane, a Myc-tagged anti-mCherry nanobody (LaM4)^39^, a Notch-core trans-membrane domain and a transcriptional Tet activator domain; The system triggers the downstream gene expression of *dLuc* in response to the presentation of the mCherry antigen on the cell membrane of sender cells (Fig. 2A). First, we used the original synNotch ligand (synNL), consisting of a signal peptide, mCherry and a trans-membrane domain of human PDGFRB (Fig. 2B). To mimic protein turnover of natural Delta ligands^40^, we added a PEST degradation domain to the carboxyl-terminus end of the synNL. We applied cyclic blue light-stimulation with a 4-h period to co-cultures of sender and receiver cells. In time-lapse bio-luminescence recordings, luminescence signals from the synNR-receiver cells started to to increase at approximately 4 h after starting light stimulation (Fig. 2C); this response is much slower than for Notch reporter cells, which showed a response at approximately 1.5 h (Extended Data Fig. 3C). The luminescence signals from synNR-receiver cells later reached a steady state without any indication of a periodic pattern, indicating the failure of oscillatory signal transmission from the sender to receiver cells. These data suggest that cell-cell communication in the synNotch pathway is less dynamic than the Notch pathway.

**Fig. 2.**
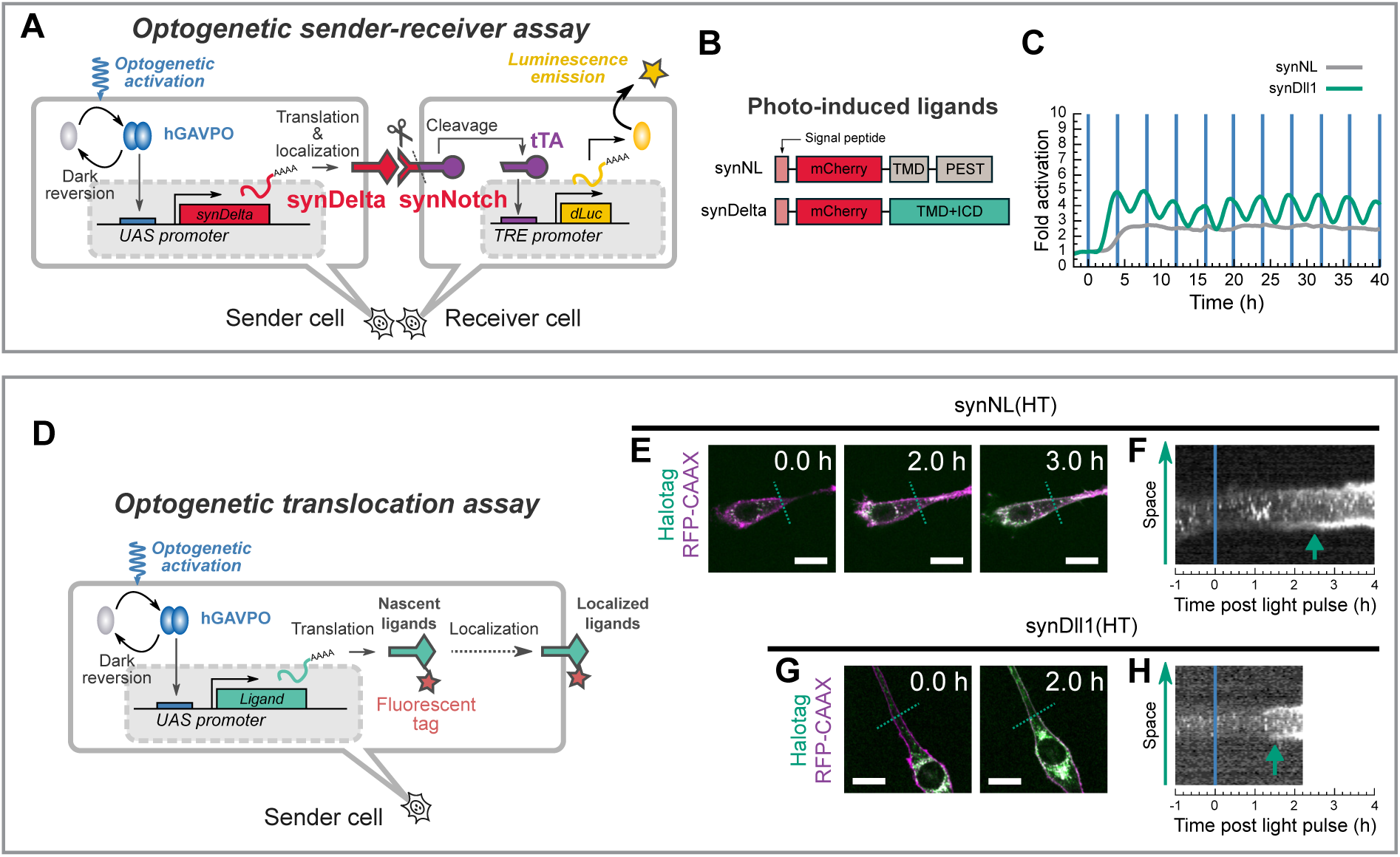
Optogenetic assays revealed dynamics of signal transmission in synthetic cell-cell signaling. (A-C) Optogenetic sender-receiver assay. (A) Schematic of the assay. (B) Schematic molecular structures of synthetic ligands and receptors used in (C). (C) Bio-luminescence signals in receiver cells illuminated with 4-h period blue light pulses. The data were acquired using a photomultiplier tube. Blue vertical lines represent the times of illumination with a 30-sec duration. (D-H) Optogenetic translocation assays. (D) Schematic of the optogenetic translocation assay. (E-H) Time-lapse imaging of translocation dynamics of photo-induced synthetic ligands labeled with fluorescent Halotag ligands. Dashed lines in (E) and (G) indicate regions used for analysis of the kymographs in (F) and (H), respectively. The labels in (E) and (G) indicate the time after a single pulse of blue-light illumination. Green arrows in (F) and (H) indicate the times of appearance of ligands at the cell membranes. Scale bars, 20 *µ*m.

We then sought to engineer synthetic cell-cell signaling that could transmit oscillatory information resembling the natural Notch pathway. We hypothesized that the failure of oscillatory signal transmission of the above synNotch (synNL-synNR) system might be due to the molecular architecture of the ligand protein. The intracellular domains of the natural Delta ligand play functional roles in clustering^41–44^, trafficking^45,46^, and force-generation^47,48^, which would also alter signal transduction dynamics. These considerations prompted us to replace the transmembrane domain of synNL with transmembrane and intracellular domains of murine Dll1, to create a new ligand, synDll1 (Fig. 2B). When we applied blue light-stimulation with a 4-h period to co-cultures of synDll1-sender and synNR-receiver cells, oscillatory signals with a 4-h period were detectable (Fig. 2C), indicating the successful transfer of oscillatory information from sender to receiver cells. Furthermore, the timing of elevation of the luminescence signals was earlier than for the sender-receiver pair of synNL and synNR.

To further characterize the signal sending process in the synthetic systems, we performed fluorescence imaging of the photo-induced ligands (Fig. 2D). We replaced the mCherry-coding sequences of synNL and synDll1 with Halotag-coding sequences and visualized the subcellular localization dynamics of the new ligands, termed synNL(HT) and synDll1(HT) respectively, using JF_646_-Halo fluorescent dyes. To identify the location of cell membranes, we used a membrane-localized red fluorescent protein (RFP); this protein is a fusion product of mScarlet-I and a farnesylation motif of HRAS (RFP-CAAX). We found that the synNL(HT) proteins appeared on the membranes at about 2.5-3.0 h after a pulse of blue light illumination, whereas the synDll1(HT) proteins appeared at about 1.2 h after the light-pulse (Fig. 2, E to H). These results suggest that the newly engineered ligand synDll1 is transported to the cell membrane more rapidly than synNL, and that membrane traffic contributes to the distinct dynamic patterns in cell-cell communication.

### 3 Engineering synthetic cell-cell signaling with tunable transmission speed

We next asked whether intracellular domains of other ligands could alter the dynamic patterns in cell-cell communication (Fig. 3A). The transmembrane and intracellular domains of murine epidermal growth factor (EGF) were used to construct a synthetic ligand, termed synEGF, in photo-induced sender cells. Optogenetic sender-receiver assays showed that a 4-h period light stimulation triggered the oscillatory responses in receiver cells (Fig. 3B); the timing of the signal-transmission was slightly delayed compared to synDll1. The use of the ligands murine Delta-like4 (Dll4) and zebrafish DeltaC in this assay showed that it was possible to generate dynamic communication channels with distinct temporal schedules (Fig. 3B and C). We also analyzed the effect of truncation of the intracellular domain of synDll1 and found that the shorter the length of the truncation mutant, the longer the delay in cell-cell communication (Extended Data Fig. 4). These data suggest that the intracellular domains of ligands have functional roles in determining the temporal schedules of cell-cell interactions.

**Fig. 3.**
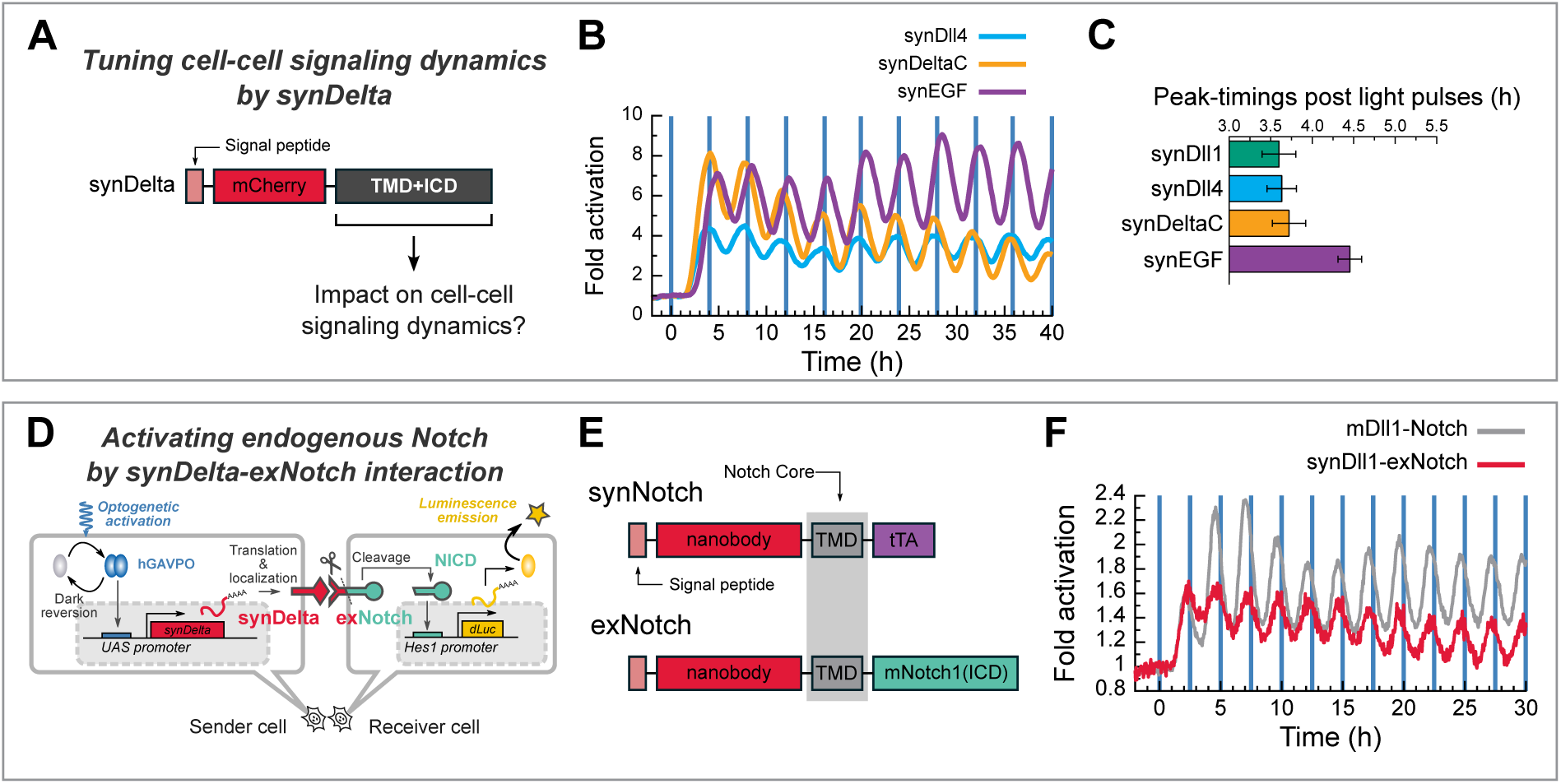
Engineering a toolbox for synthetic cell-cell signaling. (A-C) The impact of intracellular domains of ligands on cell-cell signal transmission. (B) Population signaling traces of TRE promoter activity in the presence of cyclic light illumination with a 4-h period. The data were acquired using a photomultiplier tube. Blue vertical lines represent the times of illumination of a 30-sec duration. (C) Comparison of peak timings between different synthetic ligands. n = 10 pairs of comparative peaks. (D-F) Converting synthetic signal-inputs to natural Notch signaling outputs with exNotch receptors. (D) Schematic of optogenetic sender-receiver assay that test the exNotch receptor in receiver cells that carry the exNotch receptor and Hes1 promoter luciferase reporter. (E) Schematic structures of the synNotch and exNotch receptor. (F) Population signaling traces of Hes1 promoter activity in the presence of cyclic light illumination with a 2.5-h period. The data were acquired using a photomultiplier tube. Blue vertical lines represent the times of illumination of a 30-sec duration. The data of natural Delta-Notch pairs were derived from a previous publication^36^.

The synNotch system is sufficiently flexible to allow exchange of the extracellular domains of the receptors. We replaced the LaM4 (*K_d_* of 0.18 nM) extracellular domain of the synNR with the LaM8 nanobody, which has a lower affinity (*K_d_*of 63 nM) for mCherry^39^. Optogenetic sender-receiver assays showed that this exchange slightly delayed the timing of receiver responses (Extended Data Fig. 5). In a similar manner, we also replaced the extracellular domains of synNL and synNR with EBFP and LaG17 (anti-GFP nanobody, *K_d_* of 50 nM)^39^ to create the novel ligand synNL(EBFP) and receptor synNR(LaG17), respectively. Sender cells with the photo-inducible synNL(EBFP) transmitted oscillatory information to synNR(LaG17) receivers with moderate amplitudes (Extended Data Fig. 6, A and B). Further replacement of the transmembrane and intracellular domains of synNL(EBFP) with those from mouse Dll1 to generate the ligand synDll1(EBFP), resulted in acceleration of the speed of signal transduction between sender and receiver cells. However, the fold change or the amplitude of receiver responses did not outperform that of the synDll1-synNR channel using mCherry-LaM4 pairs (Extended Data Fig. 6, B and C). Taken together, these results suggest that engineering of intracellular domains of ligands enabled tunable changes to transmission speed and amplitude of synthetic cell-cell communications through use of new ligands that we term the synDelta series.

### 4 Rewiring natural genetic circuits of the segmentation clock to program synchronized oscillation with synthetic cell-cell signaling

Connecting outputs of synthetic receptors to the gene regulatory network, such as natural Notch signaling, of the segmentation clock may be able to control synchronized *Hes7* oscillation in miPSM cells. We created a synthetic receptor, termed exNotch, which could output Notch signaling upon synthetic signaling cues, by <u>ex</u>changing the <u>ex</u>tracellular domain of mouse Notch1 with Myc-tagged anti-mCherry LaM4 nanobody fused to the signal peptide derived from human CD8*α* (Fig. 3, D and E). When the membrane-anchored mCherry antigen on sender cells interacts with an exNotch receptor on receiver cells, the intracellular domain of exNotch is cleaved and transactivates downstream genes of Notch targets, such as *Hes1*. We constructed a receiver cell that carried an expression cassette of *exNotch* and an *Hes1* promoter luciferase reporter. Optogenetic sender-receiver assays with light-inducible synDll1 sender cells showed that the temporal patterns of cell-cell signal transmission between synDelta-exNotch resembled those of natural Delta-Notch signaling (Fig. 3F).

Having established the miPSM and synDelta-exNotch systems, we asked whether engineered synthetic cell-cell signaling could function to control somitogenesis in miPSM cells. The periodic formation of somites is supported by the synchronized oscillation of Delta-Notch signaling in the PSM cell population^10,11^. We therefore designed a synthetic genetic circuit based on the natural circuit of zebrafish PSM cells; in this synthetic circuit, we rewired the molecular network of the *Hes7* oscillator in *Dll1*-KO miPSM cells by driving *synDelta* expression under the control of pHes7 and constitutively expressing *exNotch* (Fig. 4A).

**Fig. 4.**
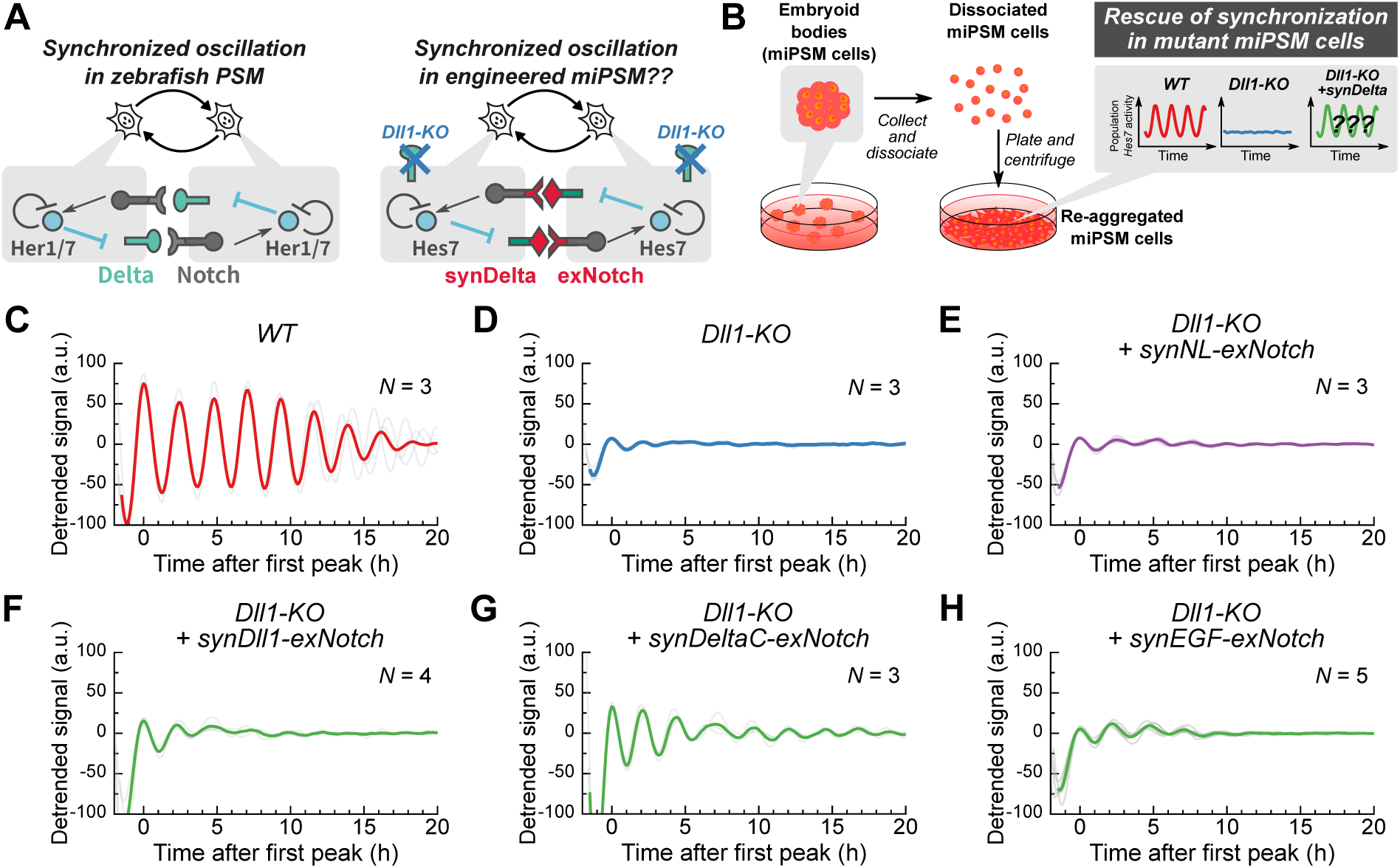
Synchronized oscillation of the segmentation clock with synthetic cell-cell signaling. (A) Schematic of the molecular networks in the segmentation clocks of zebrafish and of *in vitro* induced PSM cells that lack the natural Delta-Notch pathway but carry synthetic pathways. (B) Schematic of re-aggregation assays to test the functional capability of the population of oscillators for synchronized oscillation. (C-H) Re-aggregation assay of *Dll1*-KO induced PSM cells that carry pHes7-YFP reporter. *N* denotes number of independent experiments.

We used a re-aggregation assay to determine whether synDelta-exNotch signaling could rescue the collective behavior of *Dll1*-KO miPSM cells^15–17^ (Fig. 4B). EBs from miPSM cells were dissociated into single cells, then mixed, and re-aggregated by centrifugation. A re-aggregation assay of wild-type miPSM cells showed synchronized oscillation of pHes7 activity at the population level that was persistent for more than 20 hours after re-aggregation (Fig. 4C). By contrast, weaker pHes7 signals and rapidly dampened *Hes7* oscillations were observed in a population of re-aggregated *Dll1*-KO miPSM cells (Fig. 4D), suggesting a lack of synchronization in *Hes7* oscillation. We then integrated *synDelta-exNotch* expression cassettes into *Dll1*-KO mESCs (supplementary text) and produced miPSM cells for the reaggregation assays. We found prolonged oscillation at the population level in re-aggregated miPSM cells compared with *Dll1*-KO miPSM cells, although the amplitudes were smaller and dampened earlier than in wild-type miPSM cells (Fig. 4E-H). These results suggest that the *synDelta-exNotch* pathways partially rescued synchronized oscillation of Hes7 in *Dll1*-KO miPSM cells.

Finally, we tested whether the synDelta-exNotch pathway could program somitogenesis in organoids derived from *Dll1*-KO miPSM cells (Fig. 5A). Time-lapse microscopy of miPSM-organoids derived from *Dll1*-KO mESCs with *synDeltaC-exNotch* showed oscillatory expression of *Hes7* and propagation of waves with higher amplitudes than in organoids with *Dll1*-KO or *Dll1*-KO with *synNL-exNotch* (Fig. 5B). The number of *Hes7* waves in *Dll1*-KO miPSM tissues with *synDeltaC-exNotch* was comparable to that of wild-type miPSM tissues, while *Dll1*-KO and *Dll1*-KO with *synNL-exNotch* displayed a smaller number of waves than the wild-type (Fig. 5C, Extended Data Fig. 7). *In situ* HCR analysis indicated that expression of the somite-marker *Uncx4.1*, which was undetectable in *Dll1*-KO, was present in the rescued organoids, although the spatial pattern was disrupted compared to the wild-type (Extended Data Fig. 8). Taken together, these data suggest that engineered synthetic signaling using synDeltaC-exNotch can program the segmentation clock and somitogenesis in organoids of PSM-like tissues derived from mESCs.

**Fig. 5.**
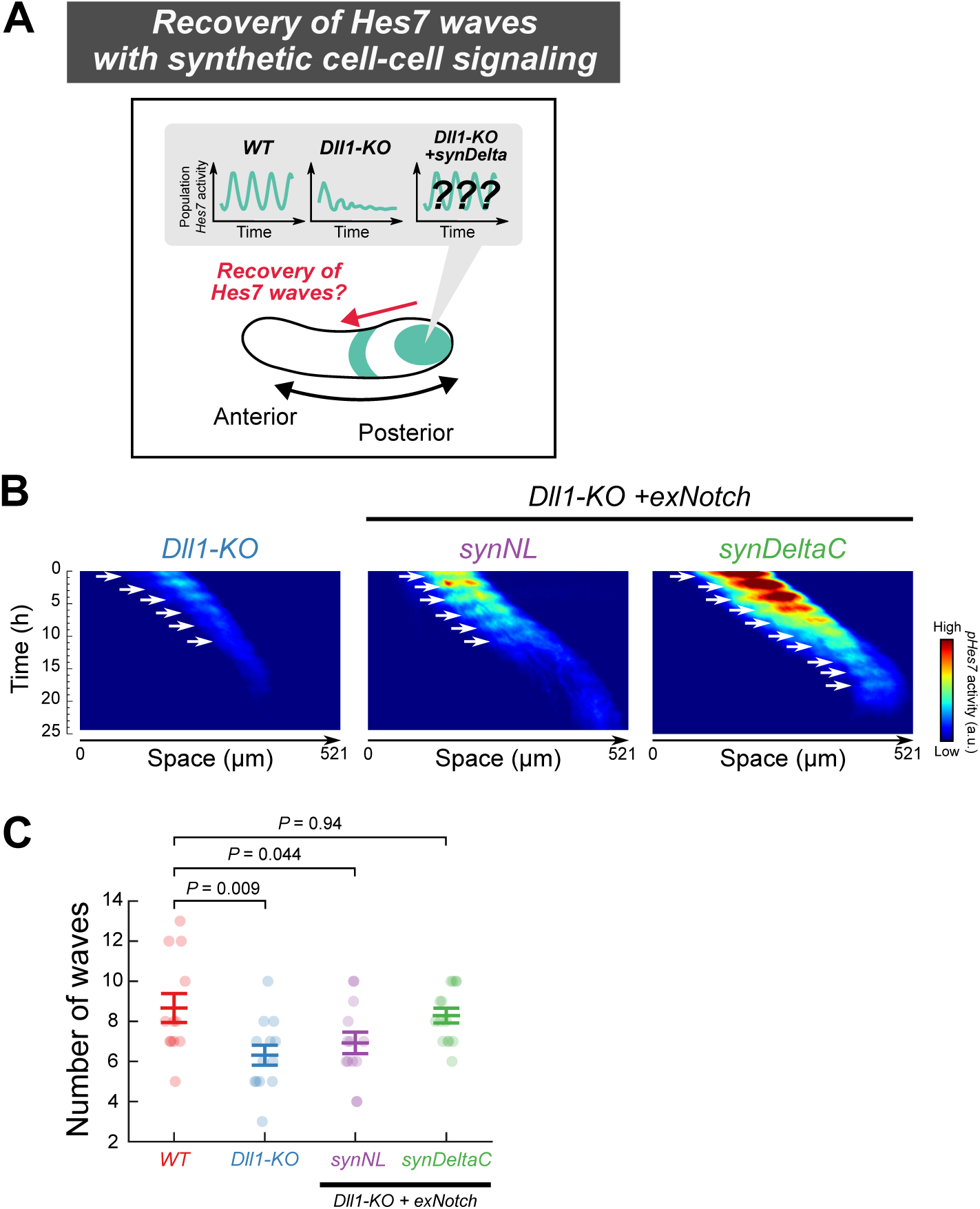
Segmentation clock waves in organoids of miPSM using synthetic cell-cell signaling. (A) Schematic of rescue experiments. (B) Time-lapse fluorescence imaging of the pHes7-Ub(G76V)-NLSA-chilles reporter in organoids of *Dll1*-KO miPSM with or without synDelta-exNotch. White arrows represent *Hes7* waves. (C) Numbers of Hes7 waves in organoid cultures of miPSM cells. *P*-values are from a one-way analysis of variance (ANOVA) with Dunnett’s post hoc test comparisons. Data are shown as the mean ± SEM; from left to right, the number of measured samples was *N* = 12, 13, 12, and 15.

## Discussion

The current paradigm for the synchrony of the segmentation clock proposes a model in which PSM cells send and receive oscillatory information through Delta-Notch signaling^11,18^. Here, we demonstrated that synthetic cell-cell signaling by synDelta-exNotch can reconstitute synchronization of *Hes7* oscillation in Delta-Notch deficient PSM cells. The ligand-receptor pair of synDeltaC-exNotch, which allows oscillatory cell-cell communication, produced a similar number of oscillatory waves in miPSM organoids as in wild-type organoids. Conversely, non-oscillatory cell-cell coupling with synNL-exNotch failed to rescue the normal pattern of *Hes7* oscillations. Our results provide the first evidence that oscillatory cell-cell coupling is sufficient for synchronized *Hes7* oscillations in PSM cells.

The extracellular domains of the synDelta ligands and the exNotch receptor used in this study lack EGF repeats that are essential in post-translational modification, such as glycosylation. Thus, the synthetic circuits of synDelta-exNotch in *Dll1-KO* miPSM cells reconstruct a simplified version of the signaling network for coupling PSM cells compared to the wild-type. A number of studies have highlighted that core ligand-receptor pairs of Delta-Notch signaling and additional factors, including *Lfng* and *Dll3* in mouse and *DeltaD* in zebrafish, are required for the synchronization of the segmentation clock^11^. Expression of *Lfng* in PSM cells exhibits oscillatory patterns in the mouse but not in zebrafish, indicating that the regulatory mechanism of Notch signal modifiers is not evolutionarily conserved. The simplified design of our synthetic circuit can provide insights into a prototype of the segmentation clock in primitive vertebrates that does not rely on the modification factors of Notch signaling.

Our study developed a method for analyzing the impact of oscillatory cell-cell signaling on stem/progenitor cell behavior. Oscillatory Dll1 expression, which is driven by the transcriptional repressor Hes1, is involved in maintenance of neural^40^, muscle^49^ and pancreatic^50^ stem cells. Duplication or removal of introns at the Dll1 gene locus alters the timing of Dll1 expression (cell-cell signaling delays) by lengthening or shortening time required for splicing. As a consequence, Hes1 and Dll1 oscillations are quenched, and proliferation of the stem cells is down-regulated. These observations indicate that Hes1 oscillations and the cell-cycle machinery are sensitive to the dynamic patterns of Dll1 expression; however, the detailed mechanism of this effect remains unresolved. Rewiring natural cell-cell signaling using synDelta-exNotch should enable a detailed analysis of the interplay between dynamic ligand expression and oscillatory gene expression in a range of stem cells.

Our optogenetic assays revealed an unanticipated role for the intracellular domain of the ligand protein in determining the temporal schedules of cell-cell communication. The distinct timings of cell-cell signal transduction between ligands with and without the intracellular domains are due in part to the process of ligand translocation to cell membranes after translation in sender cells. In previous studies, the intracellular domains of Dll1 and Dll4 were functionally implicated in clustering on cellular membranes of sender cells^41–44^ and, thereby, affecting the efficiency of signal-transduction to receiver cells expressing natural Notch receptors. When Delta and Notch molecules interact, the extracellular domains of the Notch receptors undergo trans-endocytosis into sender cells, which modulate both amplitude and dynamics of receiver-cell responses^42,44^; by contrast, synNotch receptors with antibody-antigen interactions bypass trans-endocytosis^51^. Ubiquitination of the intracellular domain of Dll1 is required for recycling after trans-endocytosis^52^. As the localization of ligands to cellular membranes precedes clustering, trans-endocytosis and recycling, the accelerated localization with the ligand intracellular domains described in this study is likely independent of these events. Serial truncation of the synDll1 tail show modulation of cell-cell signaling delays, but it is unclear which part of the intracellular domains generates the delay; detailed analysis of the molecular mechanisms that accelerate or decelerate the trafficking of ligands will need to be conducted in future work.

Optogenetic assays enable the measurement of the kinetic parameters of synthetic juxtacrine signaling and the development of new synthetic signaling with tunable speed in cell-cell communication. This approach will be useful for preparing and quantitatively characterizing synthetic genetic modules of various types, including autocrine and paracrine signaling, to program pattern formation in multicellular systems. A major principle of self-organized pattern formation is the reaction-diffusion mechanism by which signaling activities are amplified or attenuated in cell-autonomous reactions and propagate across cells and tissues^1^. Key parameters that determine patterns in the reaction-diffusion system include how quickly signaling activities or molecules *per se* travel and diffuse between cells. Generation of desirable patterns entails tuning of such parameters; direct measurement of the time required for secretion and incorporation of diffusible molecules in synthetic cell-cell signaling^53^ would be helpful in the rational design of synthetic genetic circuits for programmable tissue patterning.

Since the advent of synthetic biology, many studies have succeeded in reconstructing *de novo* multi-cellular behavior, although the engineered circuits were decoupled from endogenous events as much as possible^31^. Recent advances in organoid technology is demanding novel methods to guide and control tissue formation^54^, but current synthetic circuits constructed in organoids operate as artificial signaling cues unidirectional to natural circuits^55,56^ rather than as part of natural regulatory loops, limiting the range of applications. Rewiring natural gene networks with synthetic genetic circuits has provided insights into how network structures and molecular components impact on biological functions, including bacterial competence^57^, morphogen patterning^58^, cell proliferation^59^ and cellular aging^60^. In this study, we demonstrated that synthetic cell-cell signaling can rewire natural gene regulatory circuits of the segmentation clock, providing insights into the design principles of the synchronized oscillation. In the engineered circuit, inputs of endogenous signaling to the Hes7 promoter drive expression of synDeltaC, which subsequently stimulates exNotch receptors that activate native Notch pathways; the synthetic signals are incorporated into natural intercellular feedback loops of the *Hes7* transcriptional network. Thus, our work raises the possibility that rewiring natural gene regulatory networks with synthetic circuits will expand the possibilities for programming complex multicellular behaviors. Such an approach would be useful for the construction of designer organoids and tissues for future clinical applications.

## Acknowledgments

We thank Feng Zhang for sharing plasmids of eSpCas9(1.1); Wendell A. Lim for sharing plasmids of synNotch; Masaki Kawamata for advice on genome editing; Taiki Yoshimura and Rikuto Fukushima at Kyoto University for help with constructing plasmids; FACS analysis was performed at the iCeMS Analysis Center, Institute for Integrated Cell-Material Sciences (iCeMS), Kyoto University Institute for Advanced Study (KUIAS). This work was funded by Precursory Research for Embryonic Science and Technology (JPMJPR2043 and JPMJPR15P1 to A.I., JPMJPR17P1 to D.A.) from the Japan Science and Technology Agency (JST); Grant-in-Aid for Scientific Research on Innovative Areas (19H04960 and 18H04734 to A.I., 18H04726 to D.A.) from the Ministry of Education, Culture, Sports, Science, and Technology (MEXT), Japan; Grant-in-Aid for Scientific Research (B) (21H03540 and 18H03332 to A.I., 20H02875 to D.A.) from the Japan Society for the Promotion of Science (JSPS), Japan; Grant-in-Aid for Specially Promoted Research (21H04976 to R.K.) from the JSPS, Japan.

## Author contributions

A.I. and R.K. conceived the project. A.I. designed and performed the experiments.

A.I. analyzed data. D.A. synthesized the fluorescent probes. A.I. wrote the manuscript with input from all authors.

## Competing interests

The authors declare no competing interests.

**Correspondence and requests for materials** should be addressed to A.I. or R.K.

## Methods

### Plasmid construction

We used standard molecular biology techniques and Gibson assembly (NEBuilder HiFi DNA Assembly; NEB #E2621) to assemble the constructs used in this study. To construct the fluorescent *Hes7* promoter reporter, the *Hes7* promoter region (5,393-bp upstream fragment from the first codon)^61^ was fused to the coding regions of Ub(G76V) degron^62^, nuclear localization signal (NLS), the fast-maturing yellow fluorescent protein (YFP) Achilles^17^, stop-codon, *Hes7* coding sequence without start-codon and *Hes7* 3’UTR sequence. The *Hes7* promoter cassette was inserted into the Tol2 transposon vector system (a gift from the Kawakami Lab^63,64^) together with a selection cassette, in which a selection gene *Bsd* was driven by a constitutively active promoter pPGK (Extended Data Fig. 1B).

For genome editing of mouse embryonic stem cells (mESCs), we used a high-fidelity CRISPR-Cas9 nuclease, eSpCas9(1.1) (a generous gift from Feng Zhang, Addgene #71814)^65^. We constructed a plasmid vector that carries an expression cassette of eSpCas9(1.1)-P2A-EGFP driven by the CBh promoter and an expression cassette of sgRNA driven by a Glutamine-tRNA promoter^66,67^. To determine the nucleotide sequence of the sgRNA targeting murine *Dll1*, we used inDelphi^68^ and CRISPOR^69^, which suggested candidates that would maximize the probability of frame-shift mutations and minimize the probability of off-target editing, respectively. The guide sequence used for the *Dll1* knock out was: AAGGTCCTGCAGGCGCAAGG.

To generate optogenetic ligand-expression vectors, we used the previously described humanized version of GAPVO, termed hGAVPO^36,70^. We constructed all-in-one vectors that carried a UAS promoter that drives expression of ligands, genes of interest, the *Hes1* 3’UTR sequence, a constitutively active promoter pCAG, hGAVPO, the P2A peptide, the Puro selection marker, and an SV40 polyA sequence; the schematic description of this vector is pUAS-ligand-*Hes1* 3’UTR-pCAG-hGAVPO-P2A-Puro-SV40pA. All the ligands used in this study harbor an N-terminal signal sequence (METDTL-LLWVLLLWVPGSTGD; this was used in the original synNotch ligand^38^) derived from mouse immunoglobulin chains (Accession: P01658). The ligands also carried an mCherry sequence and either a transmembrane domain of human PDGFRB fused to a humanized d1PEST domain or a fragment of the transmembrane and intracellular domains derived from ligand-proteins (murine Dll1, murine Dll4, murine EGF or zebrafish DeltaC), designated synNL and synDelta, respectively. To construct transmembrane and intracellular domains of synDll1, synDll4, synEGF, and synDeltaC, we used gene fragments of mDll1(516-722), mDll4(519-686), mEGF(1027-1217), and zDeltaC(494-664), respectively. We produced nucleotide fragments of EBFP by introducing mutations of Y67H and Y146F into a parental EGFP gene in order to reduce photo-bleaching of ligands during blue light illumination; we replaced the mCherry sequence in synNL and synDll1 with the EBFP sequence to form synNL(EBFP) and synDll1(EBFP).

For optogenetic translocation assays, we constructed synthetic ligands incorporating a Halotag sequence^71^: synNL(HT) comprizes a fusion of the N-terminal signal sequence (METDTLLLWVLLLWVPGSTGD), an ALFA tag sequence (PSRLEEELRRRLTEP)^72^, a Halotag coding sequence, and a transmembrane domain of human PDGFRB fused to a humanized d1PEST domain; synDll1(HT) comprizes a fusion of the N-terminal signal sequence (METDTLLLWVLLLWVPGSTGD), an ALFA tag sequence (PSRLEEELRRRLTEP)^72^, a Halotag coding sequence, and a trans-membrane and intracellular domains of murine Dll1. For visualizing regions of cellular membrane, we inserted an internal ribosome entry site (IRES), mScarlet-I^73^, and CAAX membrane localization signal^74^ between the Puro and SV40 polyA sequences; the schematic description of the vector for optogenetic translocation assays is pUAS-ligand(HT)-*Hes1* 3’UTR-pCAG-hGAVPO-P2A-Puro-IRES-mScarlet-I-CAAX-SV40pA.

The original synNotch receptor was a fusion of a signal peptide derived from human CD8*α* (hCD8*α*(1-21)) for translocation to the cellular membrane, a Myc-tagged anti-GFP nanobody (LaG17)^39^, a mouse Notch1 minimal regulatory region (I1427 to R1752), and a transcriptional Tet activator (tTA) domain^38^. We synthesized further synNotch receptors that were responsive to mCherry by replacing the LaG17 coding sequences with either LaM4 or LaM8, which are high or low affinity anti-mCherry nanobodies, respectively^39^. Each fragment of the synNotch receptor was fused to the selection gene *Bsd* with a P2A peptide for bi-cistronic transcription driven by a constitutively active promoter pCAG. The Notch reporter was constructed by fusing mNotch1(1-1753) (mNotch1 without an intracellular domain) and tTA to the selection gene *Bsd* with a P2A peptide for bi-cistronic transcription driven by a constitutively active promoter pCAG. A *Tet* promoter reporter cassette was constructed by fusing the *Tet* promoter (pTRE3G) to the coding regions of Ub(G76V) degron^62^, a nuclear localization signal (NLS), luciferase (Luc2), a stop-codon, and the *Hes1* 3’UTR sequence. This cassette was inserted unstream of pCAG that drives either the synNotch receptor or Notch reporter expression. For synthesis of exNotch, fragments of the signal peptide (hCD8*α*(1-21)), namely Myc-tagged LaM4 and a mouse Notch1 without an extracellular domain (I1427 to K2531), were fused to the selection gene *Puro* with a P2A peptide for bi-cistronic transcription.

SynDelta modules driven by the *Hes7* promoter were constructed by fusing the *Hes7* promoter region (5,393-bp upstream fragment from the first codon)^61^ to the coding regions of synDelta (synDll1, synDeltaC, or synEGF), a stop codon, the *Hes7* coding sequence without a start codon, the *Hes7* 3’UTR sequence, and a selection cassette of Neo and SNAP-tag driven by a PGK promoter; the schematic description of vector for synDelta modules used in miPSM cells is pHes7-synDelta-*Hes7* 3’UTR-pPGK-Neo-IRES-SNAP-SV40pA. The schematic description of a vector for synNL module used in miPSM cells is pDll1-synNL-*Dll1* 3’UTR-pPGK-Neo-IRES-SNAP-SV40pA.

Nucleotide fragments of hCD8*α*(1-21), Myc-tag, mNotch1(1427-1752), tTA, and LaG17 were PCR-amplified from pHR-SFFV-LaG17-synNotch-TetRVP64 (a generous gift from Wendell A. Lim, Addgene #79128). Nucleotide fragments of EGFP and hPDGFRB(512-561) were PCR-amplified from pHR-EGFPligand (a generous gift from Wendell A. Lim, Addgene #79129). Nucleotide fragments for PCR-amplification of LaM4, LaM8, HaloTag, mScarlet-I and SNAP were synthesized by the “GeneArt strings DNA Fragments” service (Thermo Fisher Scientific) with humanized codon optimization.

### Cell culture

All mouse embryonic stem cells (mESCs) used in this study were derived from the E14tg2a cell line (RIKEN Bio Resource Center #RBRC-AES0135). mESCs were maintained in NDiff227 medium (Takara Bio #Y40002) supplemented with 3 *µ*M CHIR99021 (Cayman #13122), 1.5 *µ*M CGP77675 (Cayman #21089), 1,000 U/mL leukemia inhibitory factor (StemSure LIF; FUJIFILM Wako #199-16051), 100 U/mL of penicillin, and 100 *µ*g/mL streptomycin (Nacalai tesque #09367-34) at 37^◦^C and 5% CO_2_ in a humidified incubator; the culture conditions are formally known as the feeder-free a2i condition^75,76^.

We maintained C2C12 myoblast cells (purchased from DS Pharma Biomedical [Osaka, Japan]) at 37^◦^C and 5% CO_2_ in a humidified incubator. The growth media consisted of DMEM medium (Nacalai tesque #08458-45) supplemented with 100 U/mL of penicillin, 100 *µ*g/mL of streptomycin and 10% fetal bovine serum (FBS; Nichirei Biosciences, #175012).

### Cell line construction

To establish stable cell lines derived from E14tg2a mESCs, 0.5 *µ*g pT2A and 0.5 *µ*g pCAGGS-mT2TP plasmids were transfected using ViaFect Transfection Reagent (Promega #E4981) into cells cultured in a 12-well plate with 2 × 10^5^ cell-density. Transfected cells were expanded and drug-selected with 2.0 *µ*g/ml Puromycin (InvivoGen #ant-pr-1), or 20 *µ*g/ml Blasticidin (InvivoGen #ant-bl-1), or 300 *µ*g/ml G418 (Thermo Fisher Scientific #10131035).

The pHes7 reporter mESC line was initially generated by transfection with a reporter plasmid and Blasticidin selection. A clonal line was established by single-cell sorting into 96-well U-bottom plates (IWAKI #3870-096) using the automatic cell dispenser unit (ACDU) of a FACS (FACSAria II, BD). To generate a *Dll1*-KO mESC line, we transfected 2.5 *µ*g of a plasmid vector carrying the expression cassettes eSpCas9(1.1)-P2A-EGFP and *Dll1*-targetting sgRNA into the pHes7 reporter of mESCs using a ViaFect reagent. The transfected 1 × 10^6^ cells were then cultured in a 6-cm dish with a ViaFect reagent. Two days after transfection, EGFP-positive cells were collected using a FACSAria II. Six days after transfection, we further purified EGFP-negative cells from the FACS-sorted transduced cells and established clonal lines by single-cell sorting into 96-well U-bottom plates using the ACDU of the FAC-SAria II. We isolated genomic DNAs from each clonal cell population and performed Sanger-sequencing analysis for genotyping (Extended Data Fig. 2B).

To transduce synDelta-exNotch modules into *Dll1*-KO mESCs, we incorporated a 2:1 ratio of a synDelta and an exNotch module into plasmid mixtures of pT2A vectors for transfection. Transfected cells were expanded and drug-selected with 2.0 *µ*g/ml Puromycin and 300 *µ*g/ml G418. We labelled cells that had incorporated vectors with the expression cassette of pPGK-Neo-IRES-SNAP; transfected *Dll1*-KO mESCs were incubated in a2i medium supplemented with 500 nM fluorescent SNAP-ligand (SiR-SNAP^77^, made in-house). We also used the anti-Myc-Tag antibody (Alexa Fluor 488 conjugate, Cell Signaling Technology #2279) to detect surface expression of exNotch receptors. Cells with both positive-low Alexa Fluor 488 and positive-high SiR signals were collected, and clonal lines were established by single-cell sorting into 96-well U-bottom plates using the ACDU of the FACSAria II.

### Generation of mass-induced presomitic mesoderm (miPSM) cells and organoids from mESCs

PSM-like cells were induced using a modification of a previously described protocol^32^. We used microfabricated culture vessels (EZSPHERE 96-Well Plate; IWAKI #4860-900)^78^ for mass production of embryoid bodies (EBs). On day 0, 2.4 × 10^3^ mESCs were seeded into each well of a EZSPHERE 96-well plate in 300 *µ*L N2B27 medium supplemented with 20 ng/mL BMP4 (R&D #314-BP-010/CF), which subsequently formed ∼ 80 EBs comprising ∼300 cells in each well. The N2B27 medium is a 1:1 mixture of DMEM/F12 medium (Thermo Fisher Scientific #11039021) supplemented with 1X N-2 MAX (R&D #AR009) and Neurobasal medium (Thermo Fisher Scientific #12348017) supplemented with 1X B-27 without vitamin A (Thermo Fisher Scientific #12587010). The N2B27 medium was further supplemented with 0.1 mM 2-Mercaptoethanol (FUJIFILM-Wako #198-15781), 0.5X GlutaMax (Thermo Fisher Scientific #35050-061), 50 U/mL penicillin and 50 *µ*g/mL streptomycin. On day 2, EBs were transferred to a 15 ml tube, washed twice with DMEM/F12 medium (Nacalai tesque #05177-15) supplemented with 0.1%BSA (Thermo Fisher Scientific #15260037), 100 U/mL penicillin and 100 *µ*g/mL streptomycin, termed wash medium. The EBs were then transferred to microwell plates (AggreWell400, 24-well plate; STEMCELL Technologies #ST-34415) pre-treated by anti-adherence rinsing solution (STEMCELL Technologies #ST-07010) according to manufacture’s instruction. Alternatively, the EBs were then transferred to 6-cm or 10-cm diameter culture dishes. The transferred EBs were cultured in PSM-induction medium consisting of DMEM (Nacalai tesque #08489-45), 1X GlutaMax, 1 mM Sodium pyruvate (Thermo Fisher Scientific #11360-070), 1 mM nonessential amino acids (Thermo Fisher Scientific #11140-050), 1X B-27 without vitamin A, 100 U/mL penicillin, 100 *µ*g/mL streptomycin, 1 *µ*M CHIR99021, 100 nM LDN193189 (Sigma-Aldrich #SML0559), and a 0.5% final concentration of DMSO (FUJIFILM Wako #031-24051).

For organoid formation, we collected and transferred EBs to a 15 ml tube on day 4, and washed the organoids with wash medium. The EBs were embedded in 150 *µ*L N2B27 medium supplemented with 10% Matrigel (Corning #356231), plated on an 8-well glass bottom chamber (Lab-Tek II Chambered Coverglass; Thermo Fisher Scientific #155409PK), and incubated for 30 min at 37 ^◦^C and 5% CO_2_ for gelation. We added 300 *µ*L N2B27 medium to each well to avoid drying of the Matrigel.

### Re-aggregation assay

For re-aggregation assays, 4.0 × 10^3^ mESCs were seeded on day 0 into each well of an EZSPHERE 96-well plate in a volume of 300 *µ*L N2B27 supplemented with 20 ng/mL BMP4, which subsequently formed ∼ 80 EBs comprising ∼500 cells in each well. We generally used three wells of each EZSPHERE 96-well plate in the assay to produce ∼ 240 EBs, yielding more than 2 × 10^5^ miPSM cells at a later step of cell counting (see below). On day 2, EBs were transferred to a 15 ml tube, washed twice with wash medium. They were then transferred to AggreWell400 plates pre-treated with an anti-adherence rinsing solution, and cultured in PSM-induction medium supplemented with 3 *µ*M CHIR99021, 300 nM LDN193189, and DMSO at a final concentration of 0.5%. On day 4, we collected the EBs and transferred them to a 15 ml tube,h; they were washed twice with the wash medium to remove dead cells, and then dissociated enzymatically (TrypLE Express; Thermo Fisher Scientific #12604013) for 3 min at room temperature. Suspensions of dissociated cells were diluted with wash medium, centrifuged, and then re-suspended in wash medium and filtered using a cell-strainer (Falcon #352235). The number of cells was counted and the cells were re-centrifuged. After centrifugation, the cells were re-suspended in re-aggregation medium: PSM-induction medium supplemented with 1 *µ*M CHIR99021, 100 nM LDN193189, 10 *µ*M Y27632 (FUJIFILM Wako #036-24023), 25 ng/mL bFGF (Thermo-stable basic FGF; Thermo Fisher Scientific #PHG0369), 1 *µ*g/mL Heparin (STEMCELL Technologies #07980), 2.5 *µ*M BMS-493 (Sigma-Aldrich #B6688), and DMSO at a final concentration of 0.5%. A silicon ring with a 3 mm diameter hole was placed in each well of an 8-well glass-bottom chamber (IWAKI #5232-008), and 3 mm diameter glass surface was pre-coated with 50 μ g/mL fibronectin FUJIFILM Wako #063-05591) for 1 h at 37 ^◦^C and 5% CO_2_. After removal of pre-coated fibronectin solution, 2 × 10^5^ cells in 16 *µ*L cell-suspension were plated onto the 3 mm diameter glass surface in the hole of the silicon ring. After plating, the cells were re-aggregated by centrifugation at 400 g for 4 min at room temperature; 400 *µ*L of re-aggregation medium was added to each well to prevent drying.

### Time-lapse microscopy of miPSM cells

We obtained time-lapse movies of miPSM cells using an inverted microscope (IX83-ZDC2, Evident-Olympus) equipped with an environmental chamber (STRG-WELSX-SET, Tokai Hit) at 37 ^◦^C and 5% CO_2_ and a cooled scientific CMOS camera (ORCA-Flash4.0 V3, C13440-20CU, Hamamatsu Photonics K.K.). We used either the ×20 objective (UPLSAPO20X, N.A. 0.75, Evident-Olympus) for the re-aggregation assays or the ×10 objective (UPLSAPO10X, N.A. 0.4, Evident-Olympus) for organoid cultures. Fluorescence images were acquired using 2×2 binning with 16-bit resolution. Time-lapse images were collected at 6 min intervals with 2 sec exposures of the YFP channel. For fluorescence imaging, a 530 nm LED light source (LED4D, Thorlabs) and a fluorescence filter cube (YFP-2427B, Semrock) were used for excitation of the Achilles fluorescent protein. The devices were controlled using Micro-Manager software^79^ for automated image acquisition.

### Image processing and time-series analysis

Image processing and time-series analyses were used to quantify the oscillatory dynamics of pHes7-YFP signals. The images were processed using Fiji/ImageJ image-analysis software. To analyze time-lapse movies from the re-aggregation assays, temporal signals at each pixels were smoothed using a Savitzky-Golay filter (order 2, window 7) and subtracted by values of the moving average (window 31) at the population level (computed from the spatial average in the whole field of view: 665.6 *µ*m x 665.6 *µ*m area) to obtain detrended signals [gray lines shown in Fig. 4(C-H)]. Each time-series at the population level from independent experiments was further averaged to compute mean temporal traces [colored lines shown in Fig. 4(C-H)].

The time-lapse movies from the organoid assays were analyzed by smoothing the temporal traces at each pixel using a Savitzky-Golay filter (order 2, window 7) and subtracting background images of fields without a sample. Kymographs were extracted using a “Reslice” built-in function in the Fiji/ImageJ software and the Hes7 waves in the kymographs were determined manually.

### Statistical analysis

Data were analyzed statistically using the software, JASP-0.18.1^80^.

### Optogenetic sender-receiver assay

C2C12 cells were plated on 24-well plates and transfected with a 1:1 (0.5 *µ*g total amount) mixture of pCAG-mT2TP and pT2A vectors carrying either synNotch-reporter or optogenetic modules using ViaFect Transfection Reagent (Promega #E4981) according to the manufacturer’s instructions. One day later, the cells were passaged with dilution, and antibiotics were added to the culture: either 2.0 *µ*g/mL Puromycin or 20.0 *µ*g/mL Blasticidin for a week to select for transduced cells. After drug selection, the cells were collected and mixed with a 5:1 sender-receiver ratio in DMEM plus 5%FBS medium supplemented with 0.5 mM luciferin. The 1.5 × 10^5^ cells were plated in each well of black 24-well plates and used for optogenetic experiments.

Luminescence signals at the population level were recorded by a live cell monitoring system (CL24B-LIC/B, Churitsu Electric Corp.) equipped with a high-sensitivity PMT, an LED blue-light source (LEDB-SBOXH, OptoCode) and a humidified incubation system to keep cells at 37^◦^C and 5% CO_2_^36^. Photon-counting measurements were performed every 3 min with a 5 sec exposure. Light stimuli were applied every 4.0 h or 2.5 h with a 30 sec duration and an intensity of 31.2 W*/*m^2^ measured by a light meter (LI-250A, LI-COR Biosciences) unless otherwise described. Recorded traces were smoothed by a Savitzky-Golay filter (order 2, window 27) and normalized by the values at the time of light illumination (t=0) in order to calculate fold changes.

### Optogenetic translocation assay

C2C12 cells were plated on 24-well plates and transfected with a 1:1 (0.5 *µ*g total amount) mixture of pCAG-mT2TP and pT2A vectors carrying optogenetic modules of either synNL(HT) or synDll1(HT) using ViaFect Transfection Reagent (Promega #E4981) according to the manufacturer’s instructions. One day later, the cells were passaged with dilution and 2.0 *µ*g/mL Puromycin was added to the culture for a week to select transduced cells. After selection, stably transfected cells were incubated overnight under sustained blue-light illumination for induction of ligands. The culture medium was supplemented with 10 nM JF_646_-Halo (fluorescent HaloTag-ligands, made in-house)^81^ for 1 h before cell-sorting by FACS (BD FACSAria II SORP, Becton Dickinson). Cells positive for both JF_646_ and mScarlet-I signals were collected, expanded and subjected to further analysis.

We captured time-lapse movies of light-induced ligand proteins using an inverted microscope (IX83-ZDC2, Evident-Olympus) equipped with an environmental chamber at 37 ^◦^C and 5% CO_2_, the ×60 TIRF objective (UPLAPO60XOHR, N.A. 1.5, Evident-Olympus), a spinning disk confocal unit (CSU-W1, Yokogawa Electric), and a cooled qCMOS camera (ORCA-Quest, C15550-20UP, Hamamatsu Photonics K.K.). Fluorescence images were acquired using 2×2 binning with 16-bit resolution and photon number resolving mode. In fluorescence imaging, laser light sources with 555 nm and 640 nm (LDI-7, 89 North) and emission filters transmitting red (FF01-593/40, Semrock) and far-red light (FF01-680/42, Semrock) were used for visualization of fluorescent probes of mScarlet-I-CAAX^73^ and JF_646_-Halo, respectively. Time-lapse images were collected at 3 min intervals with summation of 8 images of either 100 ms exposures for 555 nm excitation or 200 ms exposures for 640 nm excitation. For optogenetic stimulation, single pulses with a 30 sec duration of 455 nm light illumination from a laser light source (LDI-7, 89 North) were applied to samples through the ×60 TIRF objective. The devices were controlled by Micro-Manager software^79^ for automated image acquisition. For image processing, we applied a spatial smoothing filter (the “Sigma Filter Plus” plug-in of Fiji/ImageJ) to the captured images.

### *In situ* hybridization chain reaction (HCR)

*In situ* HCR V3 was performed according to the previously described protocol^34,82^. We used HCR probes for *Uncx4.1* (Accession NM 013702.3; hairpin B1, Molecular Instruments) and hairpin B1 labeled with Alexa Fluor 647 (Molecular Instruments). Fluorescence images of Alexa Fluor 647 probes were visualized using an epi-fluorescence microscope (IX83-ZDC2, Evident-Olympus) equipped with a cooled scientific CMOS camera (ORCA-Flash4.0 V3, C13440-20CU, Hamamatsu Photonics K.K.), a 660-nm LED light source (LED4D, Thorlabs), and a filter-set (Cy5.5-C, Semrock).

---

**Extended Data Fig. 1.**
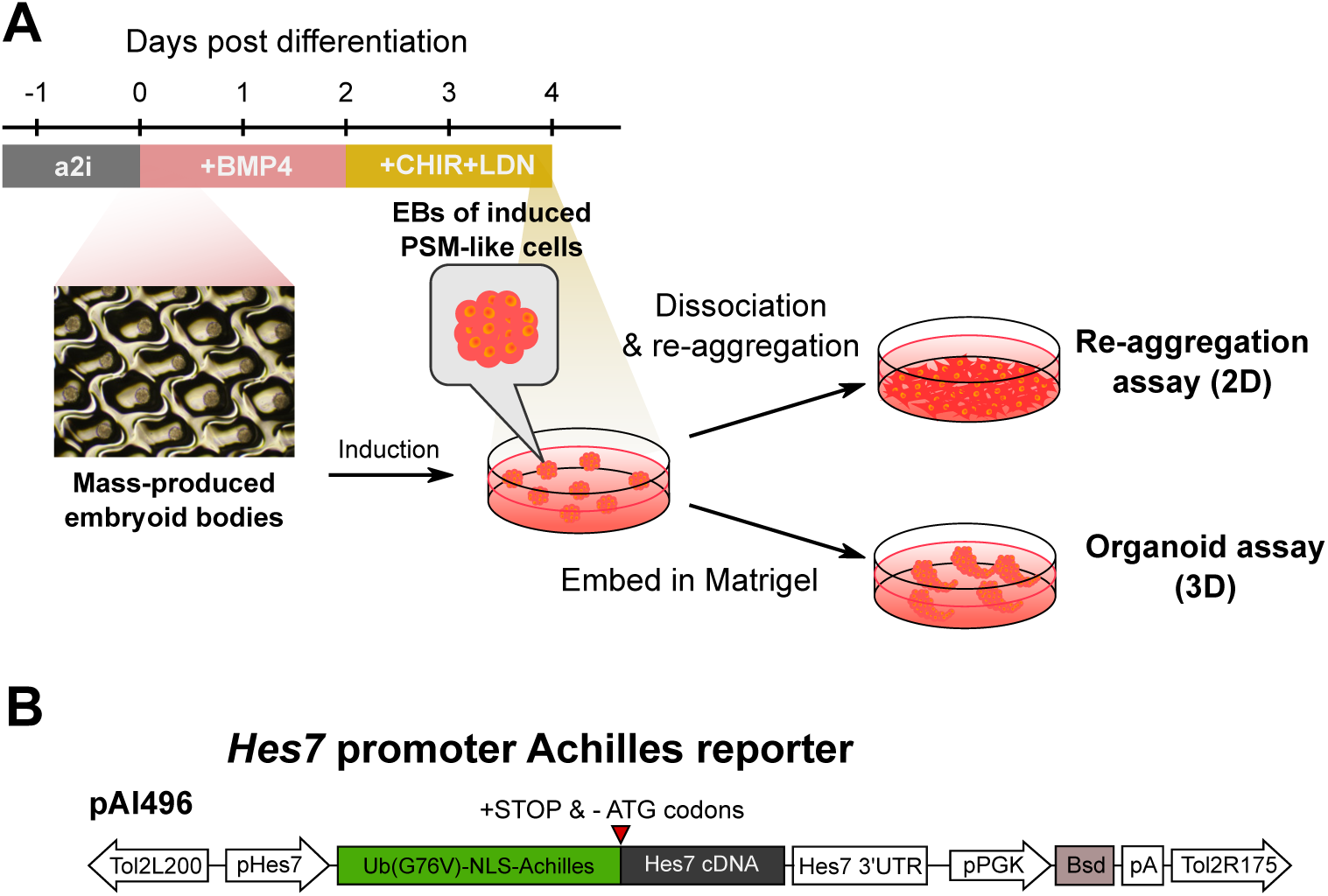
*In vitro* differentiation and visualization system for analysis of the segmentation clock. (A) Schematic of the method used in this study for mass production of PSM-like cells (termed miPSMs) from mESCs. (B) Schematic of the plasmid vector used for visualization of *Hes7* promoter activity.

**Extended Data Fig. 2.**
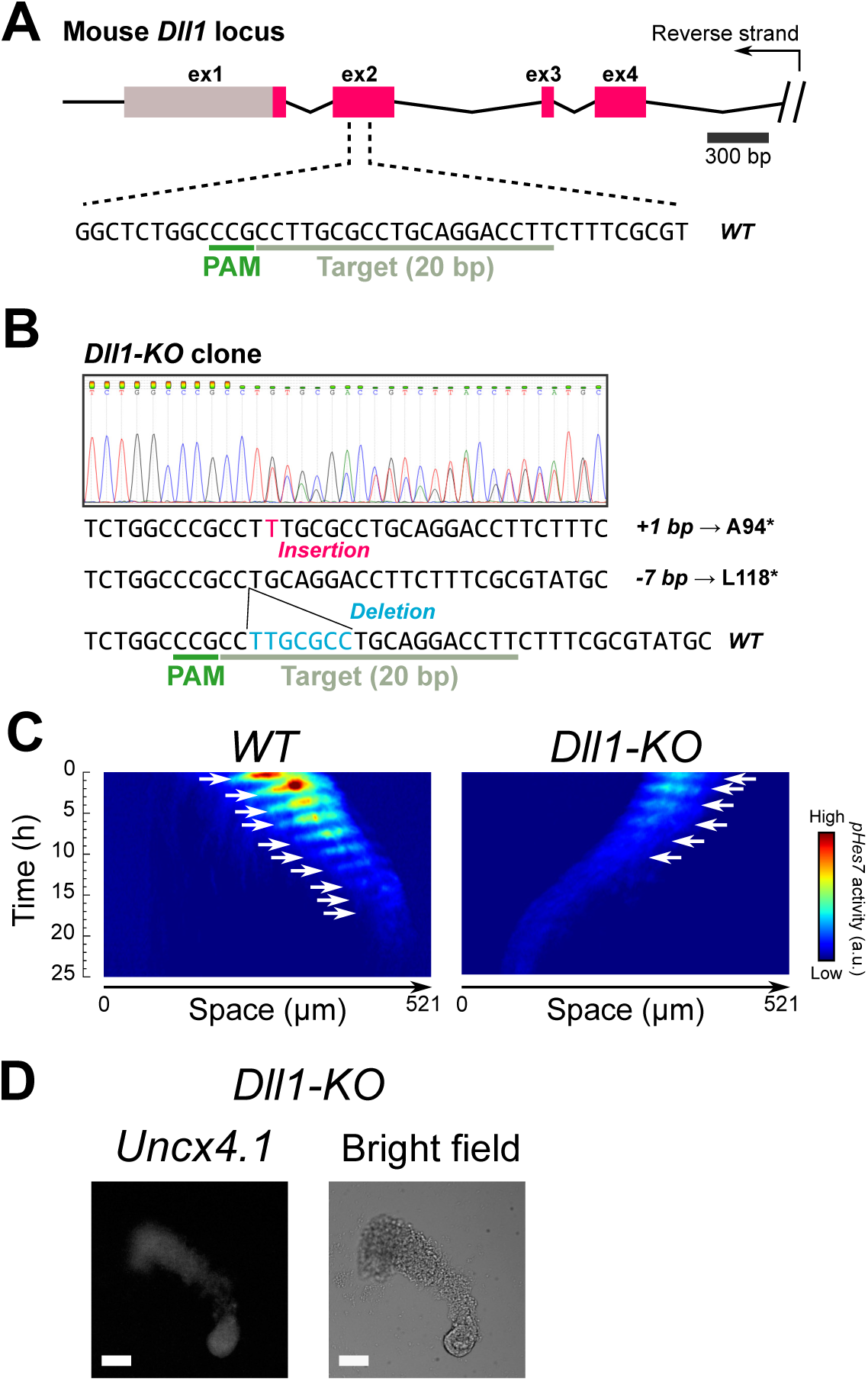
Generation and characterization of *Dll1*-KO mESCs. (A) Schematic of the method for generating the *Dll1* knockout mESC line. The positions of the sgRNAs used in this study are shown. (B) Sanger-sequence analysis of the *Dll1*-KO mESC clone used in this study showing insertion and deletion mutations induced by CRISPR-Cas9. (C) Kymographs of time-lapse fluorescence-imaging of pHes7-Ub(G76V)-NLS-Achilles reporter in *wildtype* (*WT*) and *Dll1*-KO miPSM. White arrows indicate Hes7 waves. (D) Hybridization chain reaction (HCR) v3.0 staining of *Uncx4.1* (somite marker gene). Scale bars, 100 *µ*m.

**Extended Data Fig. 3.**
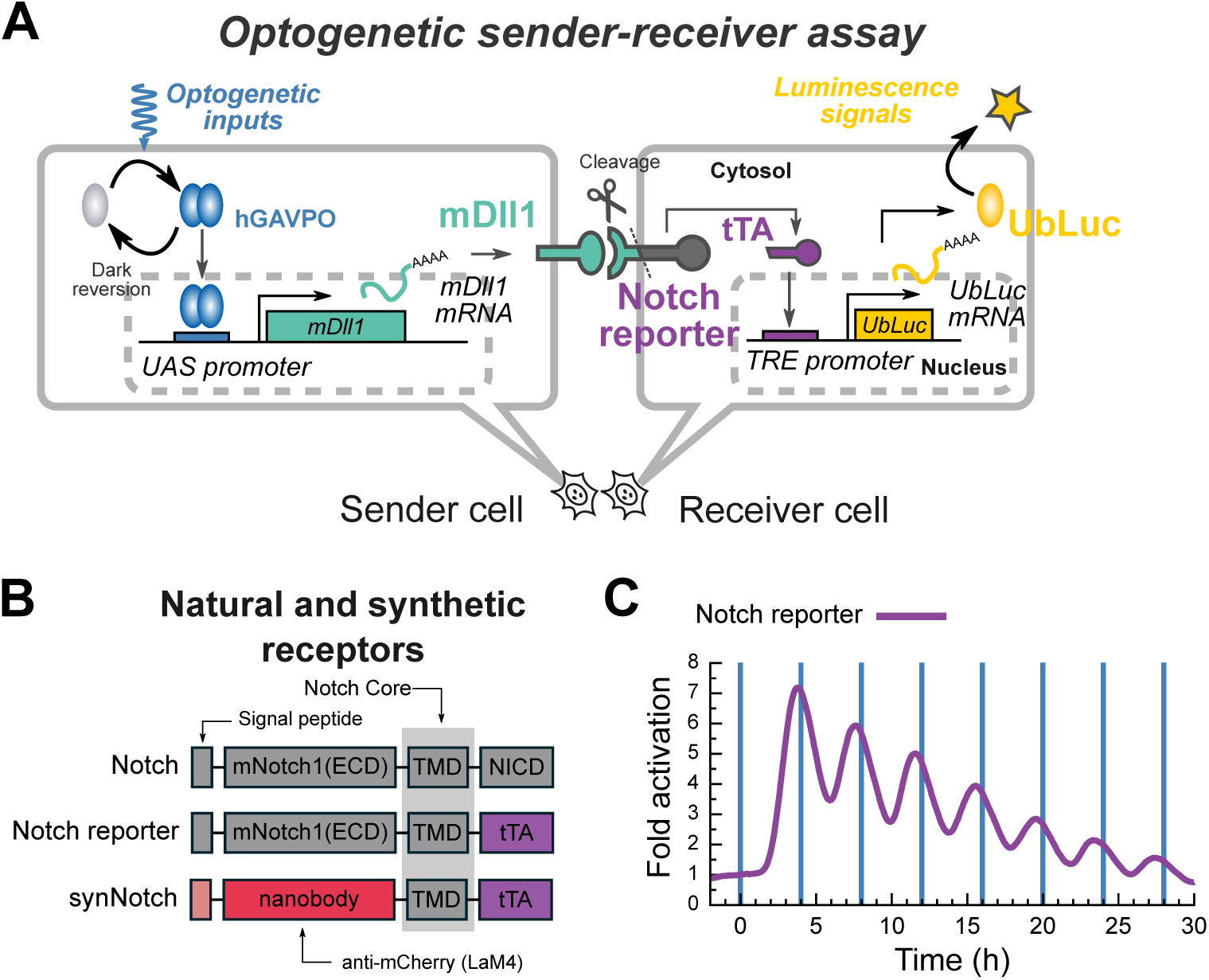
Characterization of the Notch reporter system using an optogenetic sender-receiver assay. (A) Schematic of the optogenetic sender-receiver assay for testing dynamic cell-cell signal transmission in the Notch reporter system. (B) Schematic of Notch-related receptors. (C) Population signaling traces of promoter activity of a TET responsive element (TRE) after cyclic light illumination with a 4-h period. The data were acquired using a photomultiplier tube. Blue vertical lines represent the times of illumination of a 30-sec duration.

**Extended Data Fig. 4.**
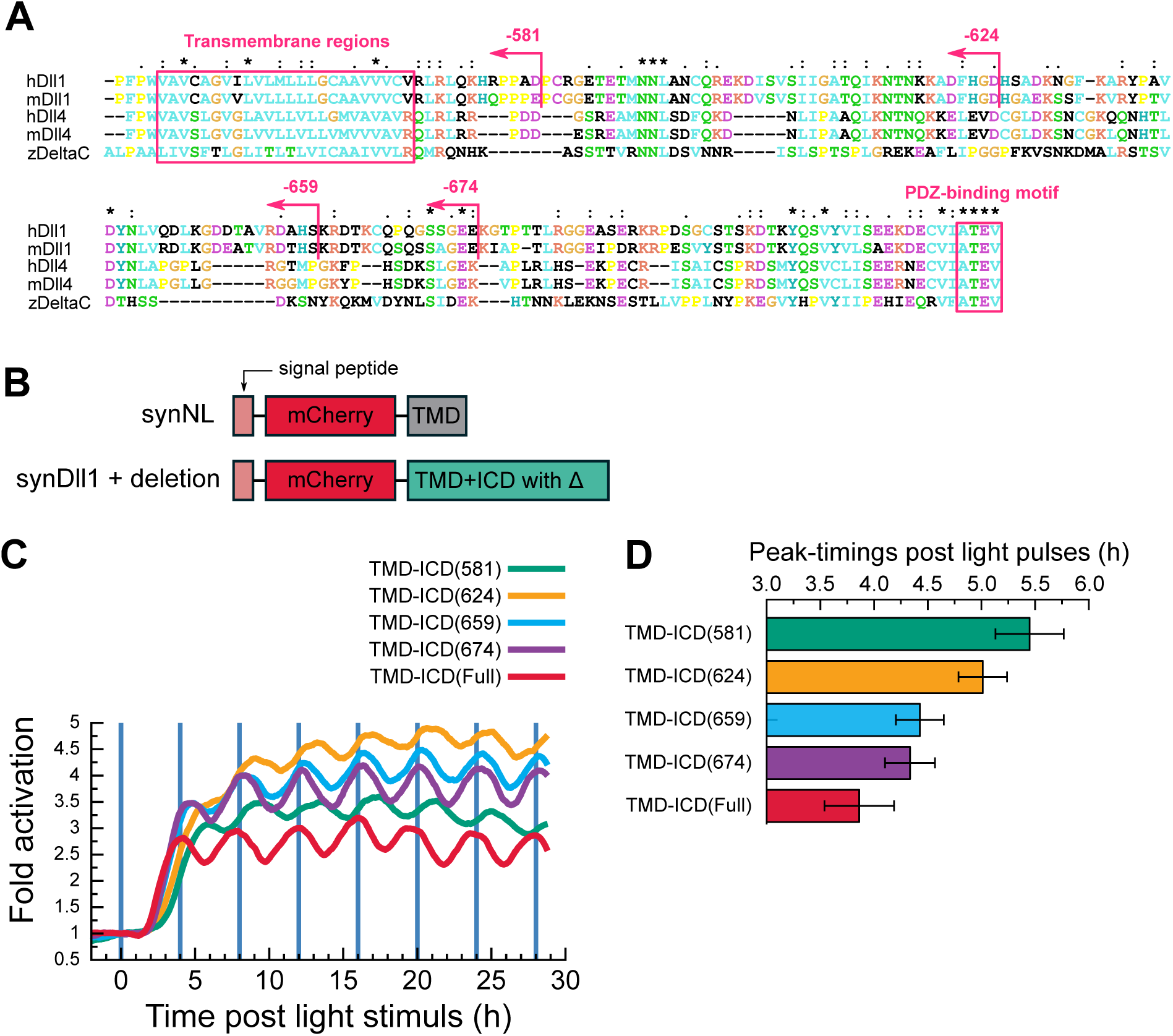
The role of intracellular domains of ligands on dynamic patterns in cell-cell communication. (A) Multiple alignments of transmembrane and intracellular domains of Delta proteins. (B) Schematic structures of the original synNotch ligand (synNL) and the newly-engineered synDll1 ligands with serial deletions in the carboxyl terminus end. (C) The optogenetic sender-receiver assay that utilizes the synNotch(LaM4) receiver cells. Population signaling traces of TRE promoter activity in the presence of cyclic light illumination with a 4-h period. The data were acquired using a photomultiplier tube. Blue vertical lines represent the times of illumination of a 30-sec duration. TMD-ICD(Full) is a synDll1 ligand containing mDll1(516-722) without the deletion. The letter X in TMD-ICD(X) denotes the positions of deletions in TMD-ICD of mDll1 and mDll1(516-X). (D) Comparison of the peak timings between different synthetic ligands. n = 7 pairs of comparative peaks.

**Extended Data Fig. 5.**
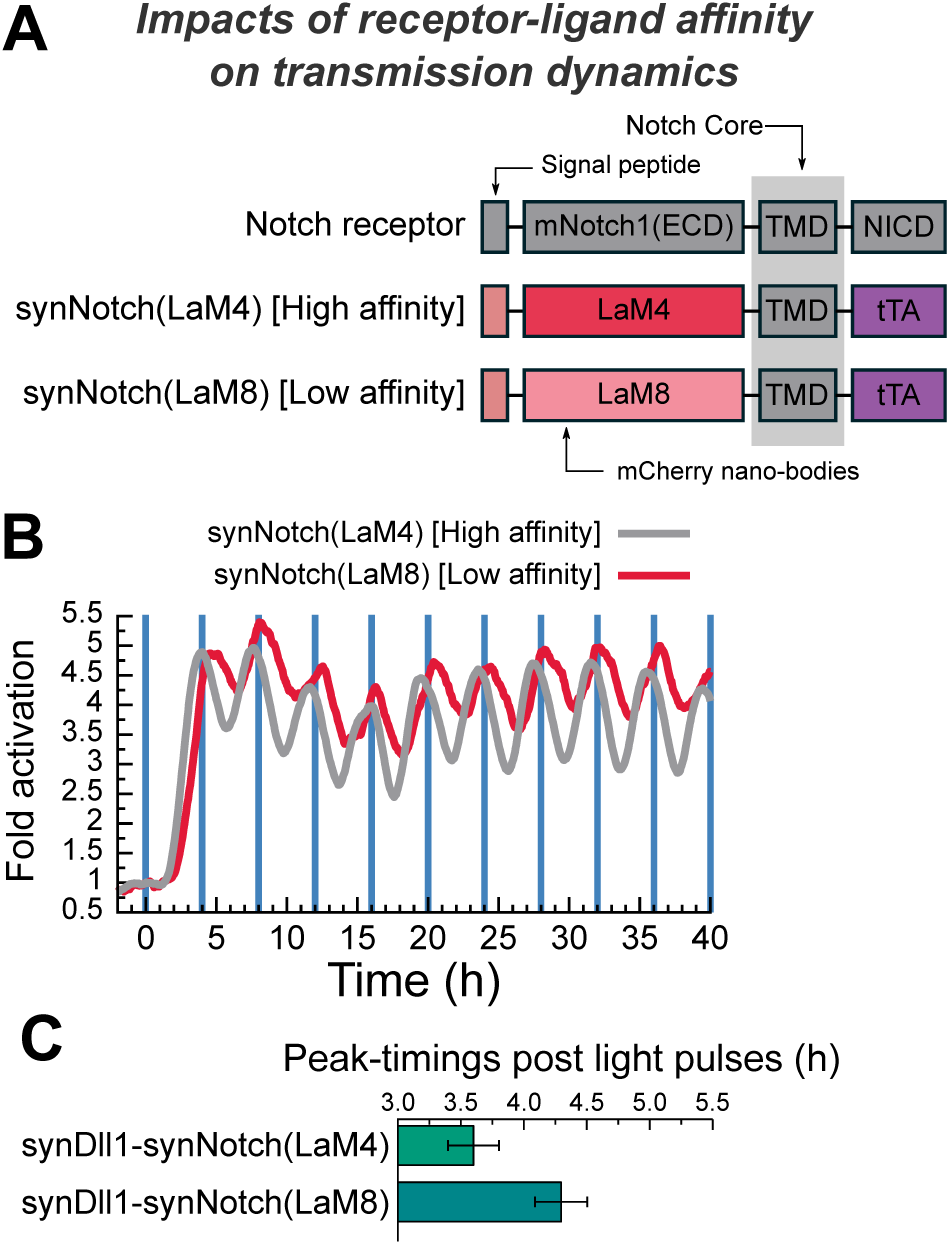
Impact of antibody-affinity of synthetic receptors on the dynamic patterns of cell-cell signal transduction. (A) Schematic structures of synNotch receptors. (B) An optogenetic sender-receiver assay that utilizes photo-inducible synDll1 sender cells. Population signaling traces of TRE promoter activity in the presence of cyclic light illumination with a 4-h period. The data were acquired using a photomultiplier tube. Blue vertical lines represent the times of illumination of a 30-sec duration. (C) Comparison of peak timings between different synthetic ligands. n = 10 pairs of comparative peaks.

**Extended Data Fig. 6.**
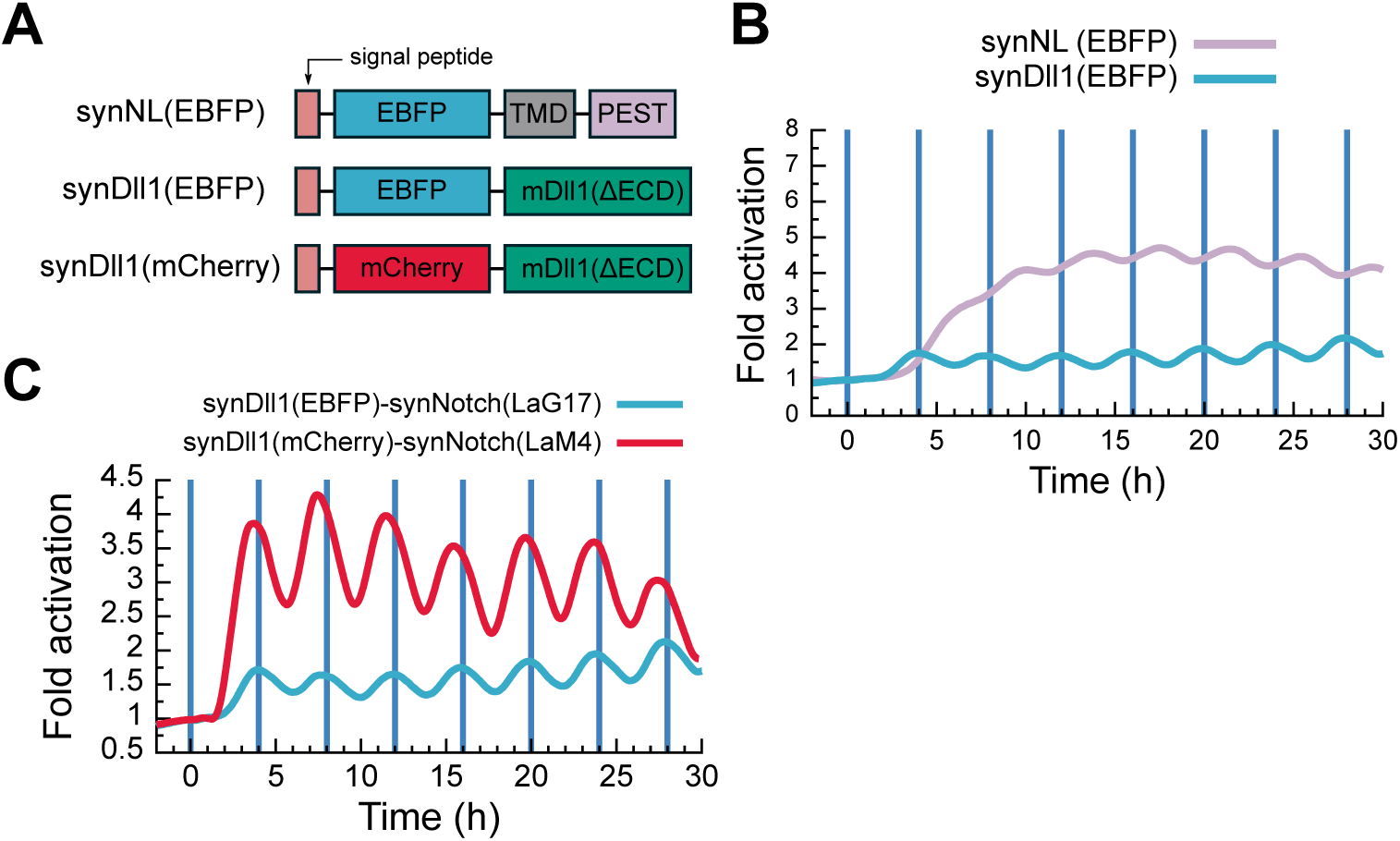
Impact of ligand-receptor pairs on synthetic cell-cell signal transduction. (A) Schematic structures of the original synNotch ligand (synNL) and newly-engineered synDll1 ligands with either mCherry or EBFP antigens. (B) An optogenetic sender-receiver assay that utilizes syn-Notch(LaG17) receiver cells. Population signaling traces of TRE promoter activity in the presence of cyclic light illumination with a 4-h period. The data were acquired using a photomultiplier tube. Blue vertical lines represent the timings of illumination of a 30-sec duration. (C) Comparison of population signaling traces between synDll1(EBFP)-synNotch(LaG17) and synDll1(mCherry)-synNotch(LaM4) pairs.

**Extended Data Fig. 7.**
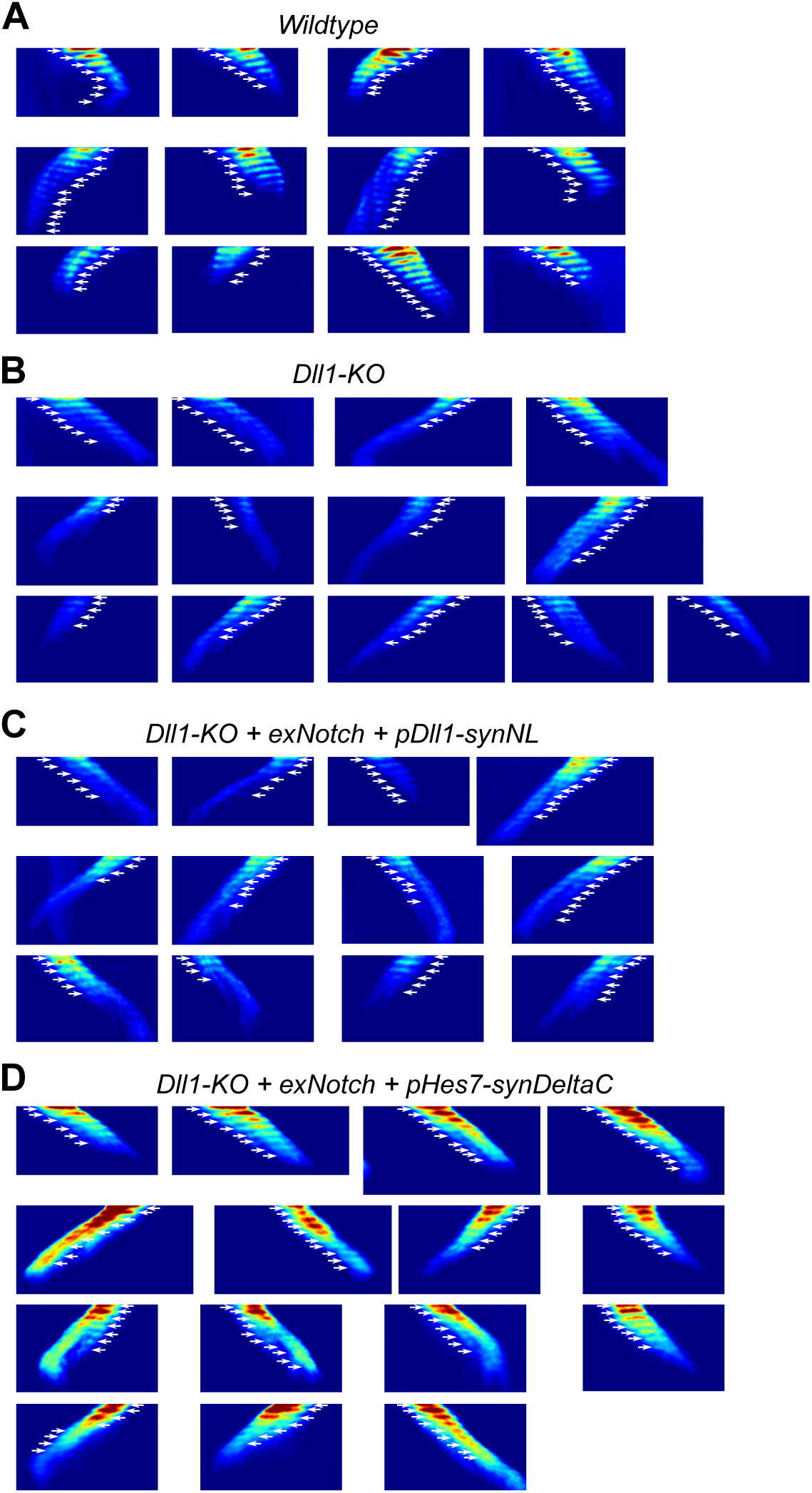
A series of kymographs from rescue experiments of somitogenesis in mass-induced PSM cells by synthetic cell-cell signaling. These data are related to Fig. 5. White arrows represent Hes7 waves.

**Extended Data Fig. 8.**
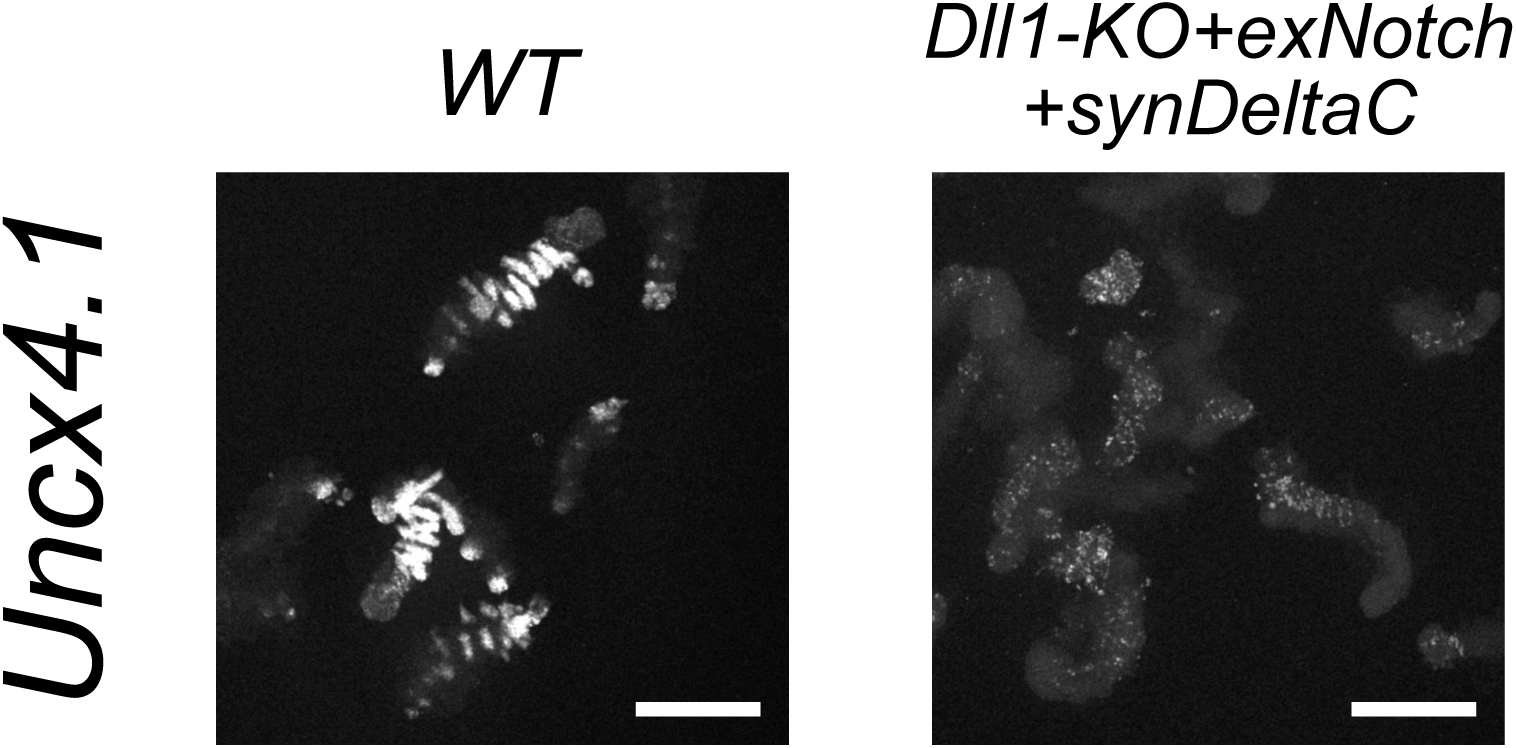
HCR staining of the somite marker gene *Uncx4.1* in wild-type (WT) and rescue (Dll1-KO + exNotch + synDeltaC) miPSM organoids. Scale bars, 400 *µ*m.

**Supplementary Table 1.** The characteristics of the plasmids used in this study.

| Plasmid name | Inserted gene | Description |
| --- | --- | --- |
| pCAGmT2TP | mT2TP | Expression vector for Tol2 transposase |
| pAI496 | Ub(G76V)-NLS-Achilles, Bsd | *Hes7* promoter reporter with destabilized YFP, Achilles |
| pAI500 | eSpCas9(1.1), EGFP | eSpCas9(1.1)-P2A-EGFP and Glu-tRNA-sgRNA for Dll1-KO |
| pAI466 | Ub(G76V)-NLS-Luc2, mNotch1(ΔICD)-tTA, Bsd | *Tet* promoter reporter with Notch-reporter receptor |
| pAI477 | Ub(G76V)-NLS-Luc2, synNotch(LaM4), Bsd | Expression vector of synNotch(LaM4) and Bsd driven by CAG promoter with Tet reporter |
| pAI531 | synNL, hGAVPO, Puro | Photo-inducible expression of synNL |
| pAI530 | synDll1, hGAVPO, Puro | Photo-inducible expression of synDll1 |
| 2807 | synNL(HT), hGAVPO, Puro, mScarlet-I | Photo-inducible expression of synNL(HT) with mScarlet-I-CAAX |
| 2809 | synDll1(HT), hGAVPO, Puro, mScarlet-I | Photo-inducible expression of synDll1(HT) with mScarlet-I-CAAX |
| 2538 | synDll4, hGAVPO, Puro | Photo-inducible expression of synDll4 |
| 2539 | synDeltaC, hGAVPO, Puro | Photo-inducible expression of synDeltaC |
| 2541 | synEGF, hGAVPO, Puro | Photo-inducible expression of synEGF |
| 2683,2686, 2688,2689 | synDll1 with deletions, hGAVPO, Puro | Photo-inducible expression of deletion mutants of synDelta-D1 |
| 2536 | Ub(G76V)-NLS-Luc2, synNotch(LaM8), Bsd | Expression vector of synNotch(LaM8) and Bsd driven by CAG promoter with Tet reporter |
| pAI465 | synNL(EBFP), hGAVPO, Puro, NLS-mCheery | Photo-inducible expression of synNL(EBFP) |
| pAI455 | synDll1(EBFP), hGAVPO, Puro, NLS-mCherry | Photo-inducible expression of synDelta-D1(EBFP) |
| pAI 442 | Ub(G76V)-NLS-Luc2, synNotch(LaG17), Bsd | Expression vector of synNotch(LaG17) and Bsd driven by CAG promoter with Tet reporter |
| pAI506 | exNotch, Puro | Expression vector of exNotch and Puro driven by CAG promoter |
| pAI534 | synDll1, Neo, SNAP | Expression vector of synDll1 driven by *Hes7* promoter |
| pAI535 | synDeltaC, Neo, SNAP | Expression vector of synDeltaC driven by *Hes7* promoter |
| pAI536 | synEGF, Neo, SNAP | Expression vector of synEGF driven by *Hes7* promoter |
| 2664 | synNL, Neo, SNAP | Expression vector of synNL driven by *Dll1* promoter |

**Supplementary Table 2.** List of the stable cell lines used in this study.

| Cell line | Plasmid | Figures | Description |
| --- | --- | --- | --- |
| 6E6 | pAI496 | 1B-E, 4C, 5C, S2C, S7A | Hes7 reporter mESC clonal line. Bsd resistant. |
| 6E6g | pAI496,<br>pAI500 (transient) | 4D, 5B-C, S2C-D, S7B | Dll1-KO mESC clonal line derived from 6E6. Bsd resistant. |
| RQ6B | pAI496, pAI506,<br>pAI534 | 4E | Rescue mESC clonal line derived from 6E6g. Bsd, Puro and Neo resistant. |
| RQ7B | pAI496, pAI506,<br>pAI535 | 4F, 5B-D, S7C | Rescue mESC clonal line derived from 6E6g. Bsd, Puro and Neo resistant. |
| RQ8A | pAI496, pAI506,<br>pAI536 | 4G | Rescue mESC clonal line derived from 6E6g. Bsd, Puro and Neo resistant. |
| RQ9F | pAI496, pAI506,<br>2664 | 4H, 5B-C, 7D | Rescue mESC clonal line derived from 6E6g. Bsd, Puro and Neo resistant. |
| D1S2 | See Ref.<sup>36</sup> | 3F, S3C | A photo-inducible Dll1 sender cell line. Reported in Ref.<sup>36</sup>. |
| CRM5 | See Ref.<sup>36</sup> | 3F | *Hes1* reporter (photo-insensitive) cell line. Reported in Ref.<sup>36</sup>. |

